# Excision of mutagenic replication-blocking lesions suppresses cancer but promotes cytotoxicity and lethality in nitrosamine-exposed mice

**DOI:** 10.1101/2021.01.12.426356

**Authors:** Jennifer E. Kay, Joshua J. Corrigan, Amanda L. Armijo, Ilana S. Nazari, Ishwar N. Kohale, Dorothea K. Torous, Svetlana L. Avlasevich, Robert G. Croy, Dushan N. Wadduwage, Sebastian E. Carrasco, Stephen D. Dertinger, Forest M. White, John M. Essigmann, Leona D. Samson, Bevin P. Engelward

## Abstract

*N*-nitrosodimethylamine (NDMA) is a DNA methylating agent that has been discovered to contaminate water, food and drugs. The alkyladenine glycosylase (AAG) removes methylated bases to initiate the base excision repair (BER) pathway. To understand how gene-environment interactions impact disease susceptibility, we studied *Aag*^−/−^ and *Aag*-overexpressing mice that harbor increased levels of either replication-blocking lesions (3-methyladenine, or 3MeA) or strand breaks (BER intermediates), respectively. Remarkably, the disease outcome switched from cancer to lethality simply by changing AAG levels. To understand the underlying basis for this observation, we integrated a suite of molecular, cellular and physiological analyses. We found that unrepaired 3MeA is somewhat toxic but highly mutagenic (promoting cancer), whereas excess strand breaks are poorly mutagenic and highly toxic (suppressing cancer and promoting lethality). We demonstrate that the levels of a single DNA repair protein tips the balance between blocks and breaks, and thus dictates the disease consequences of DNA damage.

## INTRODUCTION

DNA damaging agents lead to changes in DNA structure that can promote cancer and other diseases. Distinct types of DNA modifications can result in very different biological consequences. Two major classes of DNA damage are blocking lesions (that inhibit DNA replication) and single strand breaks (SSBs). A challenge in understanding consequences of blocks versus breaks has been that most exposures induce a complex milieu of DNA lesions. To overcome this barrier, we manipulated the initial step in base excision repair (BER) to favor either excess blocking lesions or excess SSBs. Specifically, the mammalian alkyladenine DNA glycosylase (AAG; also known as MPG or ANPG) initiates BER by removing the damaged base to create an abasic site, which is then cleaved by AP endonuclease (APE1). In the dominant pathway, polymerase β removes the resultant 5’ deoxyribose phosphate and inserts a single nucleotide, and the backbone is then sealed by ligation. As such, BER creates SSBs as requisite intermediates. For reviews of BER, see (Robertson et al., 2009), (Krokan and Bjoras, 2013), and (Wallace et al., 2012). By knocking out *Aag*, it is possible to study the consequences of unrepaired 3-methyladenie (3MeA), a replication-blocking lesion. Conversely, by overexpressing *Aag*, the impact of elevated levels of downstream BER-intermediate SSBs can be evaluated. In this study, we find that there is a concert of responses that differ substantially between *Aag*-knockout mice (*Aag*^−/−^, (Engelward et al., 1997)) and mice containing a transgenic construct for *Aag* overexpression (*AagTg*, (Meira et al., 2009), which have ∼6x higher AAG activity in the liver compared to WT (Calvo et al., 2013)). Within days after a methylating exposure, distinct patterns of early physiological changes occur in the *Aag*^−/−^ and *AagTg* mice that correlate with either cancer (for *Aag*^−/−^) or toxicity (for *AagTg*). The results from these studies thus provide a deeper mechanistic understanding of the biological consequences of blocks versus breaks, while also pointing to composite biomarkers that can potentially be used to predict the effects of seemingly small DNA structural changes that can have large biological consequences. Importantly, results show that by simply varying the level of a single DNA repair protein, which in turn tips the balance between blocks and breaks, an exposure can have radically different disease outcomes: cancer versus death.

It is well established that AAG modulates responses of cells and animals to DNA methylating agents (Alhumaydhi et al., 2020; Allocca et al., 2017; Allocca et al., 2019; Calvo et al., 2016; Calvo et al., 2013; Engelward et al., 1996; Ensminger et al., 2014; Klapacz et al., 2010; Meira et al., 2009). However, prior studies have been almost exclusively focused on model methylating agents, for which human exposure is unlikely. Here, we have turned our attention to an environmental contaminant that is a major public health concern, *N*-nitrosodimethylamine (NDMA). NDMA is potently genotoxic and carcinogenic in animals (Dass et al., 1999; Dass et al., 1998; Nishikawa et al., 1997; Peto et al., 1991; Vesselinovitch, 1969; Weghorst et al., 1989). Its genotoxicity is dependent upon metabolic activation by CYP2E1, which produces the highly reactive methyldiazonium ion (Haggerty and Holsapple, 1990; Lee et al., 1996; Sohn et al., 1991), which reacts with DNA to create 3MeA and other DNA lesions (Beranek, 1990; Pegg and Hui, 1978; Souliotis et al., 1998; Swenberg et al., 1991). Importantly, irresponsible disposal of industrial waste has resulted in massive levels of NDMA contamination in the environment. For example, ineffective containment of more than 20 million gallons of chemical waste at the Olin Chemical Superfund Site led to NDMA contamination of the local drinking water supply. NDMA is also present in food, including processed meat, a known human carcinogen. It also contaminates many public water supplies as a result of certain water treatment processes (Kimoto, 1980; Mitch and Sedlak, 2004; Richardson, 2003; Sedlak et al., 2005). Furthermore, a recent public health crisis has emerged due to high levels of NDMA contamination in commonly used drugs taken by millions of people (Parr and Joseph, 2019; Pottegard et al., 2018; Scherf-Clavel et al., 2019; Sorgel et al., 2019; Tsutsumi et al., 2019).

While AAG has several substrates, 3MeA is among the most consequential, because its other methylated substrates are either relatively benign (*e*.*g*., 7MeG) or far less prevalent than 3MeA (*e*.*g*., 3MeG) (Strauss et al., 1975). This single aberrant methyl group inhibits replicative polymerases, possibly due to disruption of critical hydrogen bonding between the polymerase and the N3 position of adenine that are required for stabilization and extension (Doublie and Ellenberger, 1998; Larson et al., 1985; Plosky et al., 2008). Cells have two major responses for tolerating replication blocking lesions, namely, recruitment of translesion polymerases and induction of homologous recombination (HR). *In vitro* studies indicate that while lower fidelity translesion synthesis (TLS) polymerases can replicate past potentially toxic 3MeA lesions, the process is significantly mutagenic (Johnson et al., 2007; Monti et al., 2010; Plosky et al., 2008; Yoon et al., 2017). Alternatively, polymerase stalling can stimulate homologous recombination (HR) (Ait Saada et al., 2018; Engelward et al., 1996; Lambert et al., 2010; Marians, 2018; Yeeles et al., 2013). Although HR is often considered error-free, misalignments and strand slippage during HR can lead to large-scale sequence rearrangement mutations, including insertions, deletions, translocations, and loss of heterozygosity that promote cancer (Aissi-Ben Moussa et al., 2009; Anwar et al., 2017; Bishop and Schiestl, 2001; Cui et al., 2011; Gupta et al., 1997; Haigis and Dove, 2003; Kolomietz et al., 2002; Lambert et al., 2005; Lambert et al., 2010; Ogiwara et al., 2008; Pal et al., 2011; Piazza et al., 2017; Shao et al., 1999; Strout et al., 1998; Zhang et al., 2011). Given that 3MeA can induce TLS and HR, a focus of this particular study is on susceptibility of mice to 3MeA-driven point mutations and large-scale sequence rearrangements. Interestingly, the predicted effects of *Aag* overexpression are very different. High levels of AAG activity cause an increase in the levels of SSBs (Alhumaydhi et al., 2020; Allocca et al., 2019; Margulies et al., 2017; Parrish et al., 2018) that cannot be bypassed by polymerases. Indeed, SSBs promote replication fork breakdown, leading to one-ended double strand breaks (DSBs) that can both signal for cell death and trigger HR.

To learn about the progression from exposure-induced DNA damage to disease for an environmentally relevant hazardous chemical, we compared *Aag*^−/−^ and *AagTg* mice for initial DNA and tissue damage, mutations, and downstream cancer in the liver (where NDMA is metabolically activated). We found that *Aag*^−/−^ mice have a dramatic increase in mutations and cancer in the liver compared to WT mice. In contrast, mutations and cancer were significantly *reduced* in *AagTg* mice relative to WT mice. Prior studies have linked AAG-induced SSBs to cytotoxicity (Allocca et al., 2017; Allocca et al., 2019; Calvo et al., 2013; Margulies et al., 2017), and so cell death may prevent mutant cells from surviving and inducing cancer. While seemingly beneficial, the cost is that there is increased tissue damage and lethality. Interestingly, the suite of responses uncovered in *Aag*^−/−^ and *AagTg* mice also occur in WT mice, but to a lesser extent. Thus, cells with balanced BER still contend with the consequences of both unrepaired 3MeA and its downstream SSBs, but they are ultimately protected from the extreme adverse outcomes observed in mice with either too little or too much *Aag* expression. Taken together, this study points to reduced AAG activity as a risk factor for NDMA-induced liver cancer, whereas elevated AAG activity increases risk for toxicity-driven phenotypes. As people are quite variable in their AAG levels (Calvo et al., 2013; Crosbie et al., 2012; Hall et al., 1993), this study points to AAG activity levels as a potentially deciding factor for disease susceptibility for people exposed to NMDA.

## RESULTS

### NDMA is potently point mutagenic, particularly in AAG-deficient mice

While it is known that DNA damage can induce mutations that cause cancer, the relative impact of an unrepaired DNA base adduct versus its downstream intermediates on point mutations had not been explored previously. We leveraged the *gpt* delta transgenic mouse model developed by the Nohmi laboratory (Nohmi et al., 1996), which enables quantification of point mutations via heterologous gene expression in *E. coli*. Briefly, these mice contain multiple copies of the bacterial nucleotide salvage pathway guanine phosphoribosyltransferase (*gpt*) gene, which confers sensitivity to 6-thioguanine. DNA extracted from *gpt* delta mouse tissues can be packaged into phage and transformed into *E. coli*, and bacteria that receive a mutant *gpt* gene can form colonies on 6-thioguanine selection plates. The frequency of resultant colonies thus correlates to the number of mutations that occurred in the mouse.

We studied point mutations in livers from WT, *Aag*^−/−^ and *AagTg* mice exposed to 10.5 mg/kg NDMA, injected intraperitoneally in mice in a split dose at 8 (3.5 mg/kg) and 15 days old (7 mg/kg), a dosing regimen that was previously shown to cause liver and lung tumors in C57Bl6 mice (Dass et al., 1998) (Figure S1). We extracted genomic DNA from livers 10 weeks after exposure, a timepoint at which there was no histological evidence of neoplastic changes (Figure S2), thus reducing the possibility that mutations were caused by a tumor-associated mutator phenotype. We observed increased levels of hepatocellular degeneration and hypertrophy at 10 weeks in all mice treated with NDMA, but these did not differ significantly between NDMA-treated groups (Figure S2). When we analyzed point mutations, we observed a significant increase in WT mice (Figure 1A), consistent with the ability of NDMA to create point mutagenic DNA damage (Beranek, 1990; Pegg and Hui, 1978; Souliotis et al., 1998; Swenberg et al., 1991). *Aag*^−/−^ mice, on the other hand, have a greatly enhanced susceptibility to NDMA-induced mutations (Figure 1A) which is consistent with evidence that replication past 3MeA lesions via TLS polymerases leads to the insertion of incorrect nucleotides (Johnson et al., 2007; Monti et al., 2010; Plosky et al., 2008).

**Figure 1.**
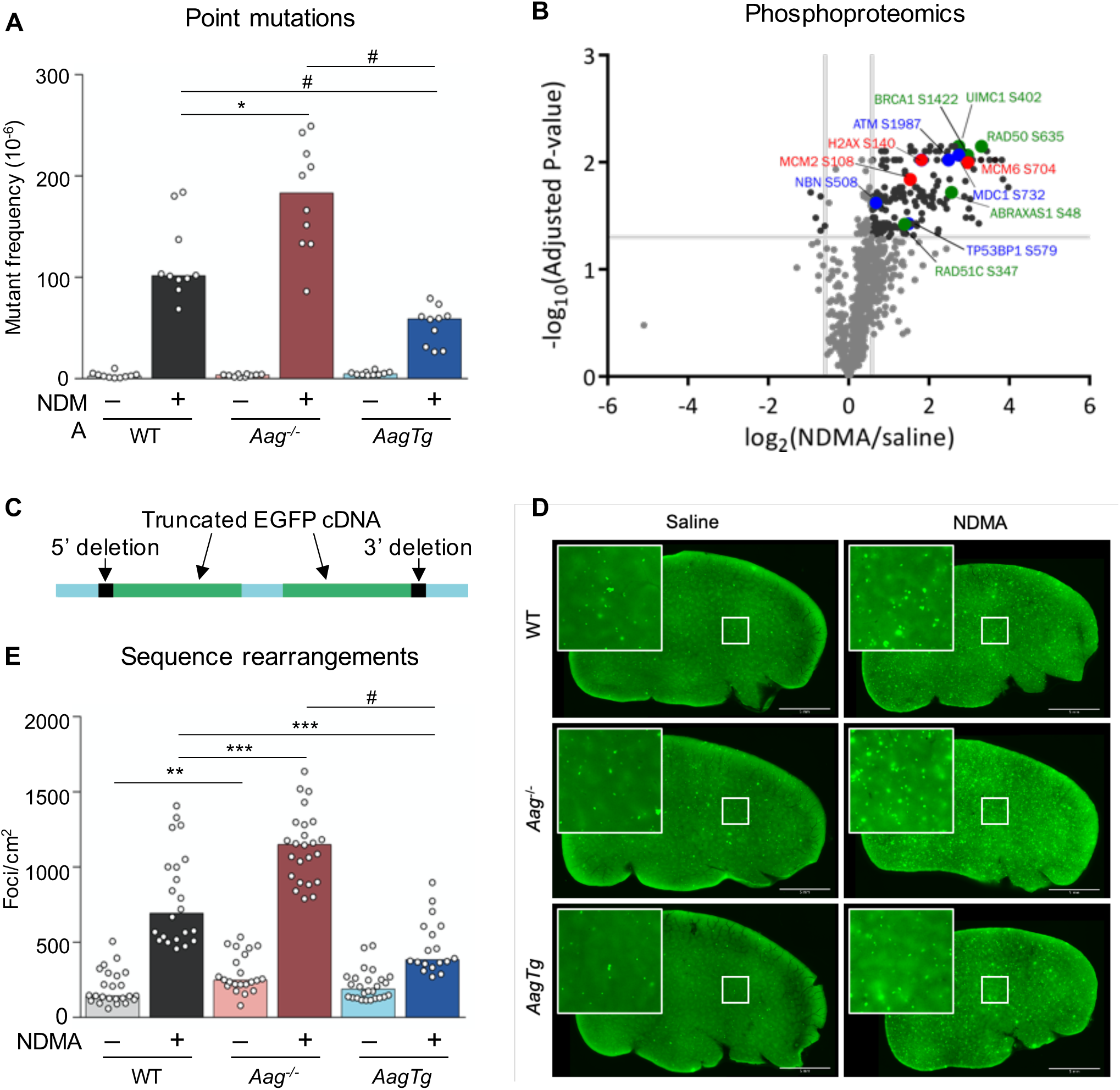
*Aag* expression levels modulate NDMA-induced point mutations, phosphoproteomic responses, and homologous recombination-driven sequence rearrangements. A) At 10 weeks after exposure, point mutations induced by NDMA were detected in the liver. Mann-Whitney *U*-test, \**p* < 0.05, ^#^*p* < 0.0001. B) NDMA treatment induces phosphorylation of proteins involved in DSB recognition (blue font), replication stress (red), and HR (green) 24h post-exposure. Volcano plot of phosphorylation sites quantified from ATM/ATR substrate motif specific (phospho-SQ/TQ) proteomic analysis in WT mouse livers treated with saline or NDMA. Log2 fold changes are relative to saline control. *P*-values were calculated based on two-tailed Student’s *t*-test and were corrected for multiple hypothesis testing based on Benjamini-Hochberg FDR correction. C) The RaDR transgenic construct consists of a direct repeat of two EGFP expression cassettes, wherein the 5’ cDNA has been truncated at the 5’ end and the 3’ cDNA has been truncated at the 3’ end. D) Representative RaDR liver images from each group, 10 months post-exposure. Scale bar = 5 mm. E) NDMA induces sequence rearrangement mutations detected 10 weeks post-exposure. Bar graphs show the median with data points representing individual animals. Mann-Whitney *U*-test, \*\**p* < 0.01, \*\*\**p* < 0.001, ^#^*p* < 0.0001.

Excision of 3MeA by AAG produces an abasic site, which can inhibit polymerases (Pages et al., 2008), and TLS bypass of abasic sites can result in point mutations (Haracska et al., 2001; Zhang et al., 2000). We therefore hypothesized that mice overexpressing *Aag* would show an increase in point mutations compared to WT mice due to increased abasic sites. However, *AagTg* mice showed a significantly *lower* frequency of point mutations than WT, and far lower than that of *Aag*^−/−^ mice (Figure 1A). This result is consistent with AAG-mediated abasic sites being quickly processed by APE1 into SSBs, which cannot be mutagenic via TLS.

### NDMA induces phosphorylation cascades indicative of HR induction

When replication forks encounter lesions that inhibit progression, such as SSBs or replication-blocking lesions, the stalled or broken fork be restored by HR. However, mis-insertion during replication fork repair can lead to large-scale sequence rearrangements that are known to drive cancer (Aissi-Ben Moussa et al., 2009; Bishop and Schiestl, 2001; Haigis and Dove, 2003; Kolomietz et al., 2002; Ogiwara et al., 2008; Pal et al., 2011; Strout et al., 1998). We therefore investigated the possibility that NDMA is recombinogenic *in vivo*. Specifically, since DNA damage induces a DNA damage response (DDR) characterized by activation of ATM and ATR kinases (Matsuoka et al., 2007), we analyzed the putative ATM/ATR substrate motif-specific phosphoproteome of control and NDMA-exposed WT mice. Analysis shows activation of DDR 24h post-exposure, consistent with stress responses to DNA damage (Figure 1B and Table S1). In particular, we observed phosphorylation of proteins associated with DSB recognition (blue font), including at the known DNA damage-regulated S732 site of MDC1 (Matsuoka et al., 2007), the ATM autophosphorylation site S1987 (Daniel et al., 2008; Pellegrini et al., 2006), and as yet uncharacterized sites on the DSB recognition proteins 53BP1 (S579) and NBN (S508). We also found that NDMA induced phosphorylation at sites associated with replication stress and DNA damage (red font), including H2AX S140 (Mazouzi et al., 2016), MCM6 S704 (Mazouzi et al., 2016), and MCM2 S108 (Charych et al., 2008; Cortez et al., 2004). Finally, we identified increased phosphorylation of HR proteins at known activation sites (green font), including RAD50 S635 (Gatei et al., 2011) and UIMC1 (aka RAP80) S402 (Matsuoka et al., 2007; Wang et al., 2007; Yang et al., 2018) as well as at uncharacterized sites on the HR proteins BRCA1 (S1422), RAD51c (S347), and ABRAXAS1 (S48). Notably, the S1422 site on mouse BRCA1 has high sequence similarity to the human BRCA1 site S1423, which is important for HR activity (Beckta et al., 2015), the G2/M checkpoint (Xu et al., 2001), and acetylation of p53 (Li et al., 2019). Given this molecular evidence that NDMA induces DNA damage, replication stress, and HR, we next turned our attention to a phenotypic readout for HR events.

### NDMA lesions repaired by AAG are more recombinogenic than their downstream repair intermediates

To more directly assess HR, we performed a functional assay for genetic sequence rearrangements 10 weeks post-exposure. The RaDR transgenic mouse model enables evaluation of the frequency of mutagenic recombination events in whole-mount tissues (Sukup-Jackson et al., 2014). The RaDR transgene (shown in Figure 1C) contains a direct repeat of truncated EGFP expression cassettes at the ubiquitously expressed *Rosa26* locus (Soriano, 1999; Sukup-Jackson et al., 2014). Upon recombination between the cassettes, full-length EGFP cDNA can be produced and expressed, creating a fluorescent signal (as shown in Figure S3). The frequency of this specific mutagenic recombination event can be readily visualized in whole-mount tissue where fluorescent foci are indicative of recombination events (Figures 1D and S3). To achieve rapid and unbiased quantification, we developed a machine learning algorithm to enumerate fluorescent foci in whole-mount liver tissue and normalize to tissue area (see STAR methods and Figure S4). Here, we demonstrate for the first time that NDMA is highly recombinogenic *in vivo* (Figure 1D and E).

As SSBs are known to cause fork breakdown, which stimulates HR, we predicted that *AagTg* mice would be highly susceptible to NDMA-induced recombination events. Unexpectedly, *AagTg* mice showed a strong *reduction* in HR events compared to WT mice (Figure 1D and E). This observation raises the possibility that damaged cells are also susceptible to cell death as well as HR. In contrast to the *AagTg* mice, *Aag*^−/−^ mice have a significant increase in susceptibility to mutagenic recombination compared to WT (Figure 1D and E). This result is consistent with prior *in vitro* studies showing that 3MeA is recombinogenic (Engelward et al., 1998; Engelward et al., 1996; Hendricks et al., 2002), and with the possibility that 3MeAs are less toxic than SSBs, allowing for more mutant cells to survive.

### DNA DSBs and chromosomal instability are modulated by AAG activity

Given that both replication-blocking lesions and SSBs can cause fork breakdown, we next asked if AAG modulates the levels of DSBs. Phosphorylation of the histone variant H2AX at Ser139 (γH2AX) occurs at sites of replication stress, most significantly in response to DSBs (Bekker-Jensen and Mailand, 2010; Chanoux et al., 2009; Scully and Xie, 2013; Stucki et al., 2005; Ward and Chen, 2001). We therefore approximated the frequency of cells that harbor an increase in DSBs 24h post-exposure by quantifying cells with five or more γH2AX foci (Figure 2A, white arrow). In WT mice, we observed a significant increase in the levels of cells with increased levels of γH2AX foci, consistent with NDMA-induced DSBs (Figure 2C). With regard to *Aag*^−/−^ mice, the high frequency of recombination events led us to predict an accordingly high frequency of DSBs.

**Figure 2.**
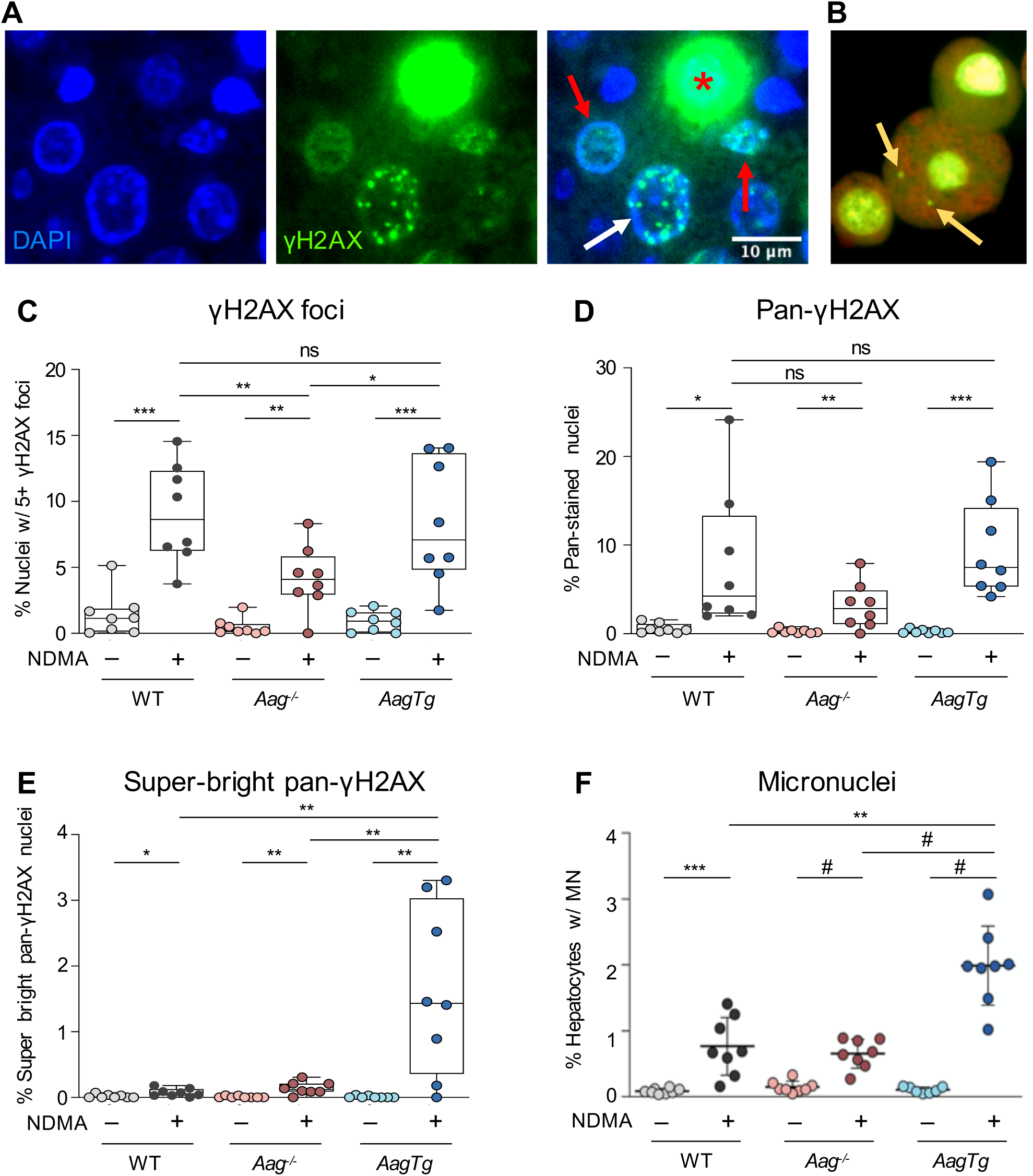
AAG levels modulate NDMA-induced phosphorylation of H2AX and micronuclei. A) Staining of γH2AX in NDMA-treated livers can appear as punctate foci (white arrow), pan-nuclear (red arrows) or super-bright pan-nuclear (red asterisk). Scale bar = 10 μm. B) Example of a hepatocyte with two micronuclei (yellow arrows). Image adapted from (Avlasevich et al., 2018) with permission from the authors and publisher. C) NDMA exposure increases the number of cells harboring an increase in punctate γH2AX foci 24h post-exposure. Nuclei containing 5 or more foci were counted as γH2AX-positive. D) NDMA increases the frequency of cells with pan-nuclear γH2AX signal 24h post-exposure. E) NDMA increases the frequency of super-bright pan-nuclear γH2AX stained cells 24h post-exposure. F) NDMA induces micronucleus formation in hepatocytes 48h post-exposure. Box plots show median with interquartile range, and whiskers show the range. Micronuclei represented as mean ± SD. For all graphs, each data point represents one mouse. All statistical comparisons were calculated by unpaired Student’s *t*-test; \**p* < 0.05, \*\**p* < 0.01, \*\*\**p* < 0.001, ^#^*p* < 0.0001.

Surprisingly, we instead observed *fewer* cells with γH2AX foci relative to WT mice (Figure 2C). These data suggest that 3MeA can induce HR via a DSB-independent mechanism, possibly via template switching. We next quantified cells with γH2AX foci in livers of *AagTg* mice (Figure 2C). We observed comparable levels of γH2AX-positive cells for *AagTg* mice compared to WT, despite the previous observation that *AagTg* mice had reduced HR-driven mutations at 10 weeks (Figure 1E). A potential explanation for these observations is that excess SSBs cause such high numbers of DSBs that cells die instead of undergoing mutagenesis. To explore this possibility, we evaluated pan-nuclear staining of γH2AX (Figure 2A, red arrows), which occurs when there is an overwhelming amount of DNA damage (Baritaud et al., 2012; Ewald et al., 2007; Moeglin et al., 2019). Although we observed similar levels of pan-nuclear staining for all genotypes (Figure 2D), during our analysis, we also observed super-bright pan-nuclear γH2AX staining, which has previously been shown occur in apoptotic cells due to cytotoxic damage in S-phase (Ewald et al., 2007; Huang et al., 2004; Moeglin et al., 2019) (Figure 2A, red asterisk). Using ImageJ (NIH), a high threshold for pixel intensity (combined with exclusion of cells with subnuclear foci) was applied to identify cells with exceptionally intense pan-nuclear staining (see STAR methods). Interestingly, *AagTg* livers showed a dramatic increase in the frequency of super-bright pan-stained nuclei, whereas WT and *Aag*-/- showed only slight increases (Figure 2E). These results indicate that *AagTg* cells have extremely high levels of DNA damage that potentiate cell death, which would be consistent with reduced mutant cells at 10 weeks.

In addition to induction of γH2AX, DSBs can also lead to the formation of micronuclei (MN) in dividing cells (an example of a hepatocyte with MN is shown in Figure 2B). We observed a significant increase in MN in NDMA-treated WT mice, a result that is consistent with DSBs during cell division (Figure 2F). (Note that division was similar among NDMA-treated groups as evidenced by Ki67; Figure S5.) Interestingly, *Aag*^−/−^ mice showed similar levels of MN induction as WT. In contrast, the *AagTg* mice had significantly higher frequencies of MN than WT, consistent with the observation that these mice are susceptible to overwhelming amounts of DSBs. Since overwhelming and persistent DNA damage can be cytotoxic, we next analyzed the frequency of cell death *in vivo*.

### NDMA causes hepatotoxicity in mice with imbalanced BER

To determine the acute hepatotoxic effects of NDMA in mice, we evaluated apoptosis and histopathological changes in livers 24 hours after the second NDMA injection. Consistent with induction of cytotoxic DNA damage, we observed an increase in apoptotic events in all NDMA-treated livers, with significantly more apoptosis in *Aag*^−/−^ compared to WT and the highest degree in *AagTg* livers (Figure 3A). Histologically, all groups of NDMA-treated mice had changes consistent with hepatocellular necrosis and degeneration (Figure 3B and C), which were often noted in centrilobular regions (Figure S6B, D, F, H, J, and L). Remarkably, *AagTg* livers exhibited significantly higher histological scores of centrilobular necrosis compared to both WT and *Aag*^−/−^ (Figure 3B). The centrilobular lesions in *AagTg* livers contained multiple necrotic hepatocytes, characterized by increased cytoplasmic eosinophilia and nuclear fragmentation, which were surrounded by vacuolated eosinophilic hepatocytes (degeneration) and low numbers of leukocyte infiltrates at the periphery (Figure S6J and L). *Aag*^−/−^ mice had increased hepatocellular necrosis and degeneration compared to WT mice (Figure 3B and C), with multifocal areas of hepatocellular degeneration mixed with an increased number of necrotic hepatocytes and inflammatory cell infiltrates (Figure S6F and H). WT livers showed multifocal areas of mild hepatocellular degeneration mixed with foci of single cell necrosis in centrilobular zones (Figure S6B and D). Together, these lesions were consistent with previous studies in laboratory rodents exposed to NDMA (Barnes and Magee, 1954; George et al., 2019; Tolba et al., 2015).

**Figure 3.**
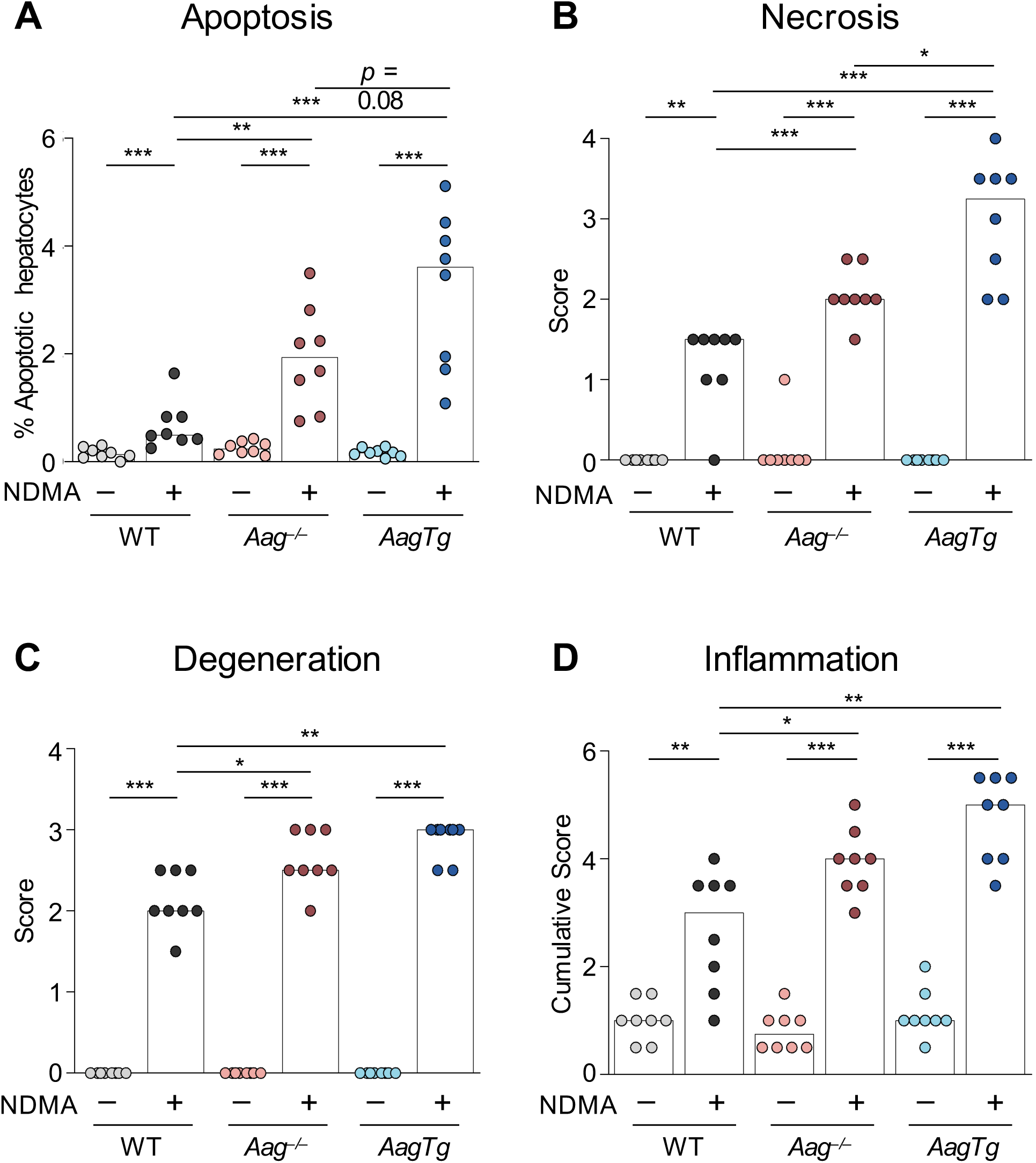
Imbalanced BER exacerbates NDMA-induced cytotoxic liver damage 24h post-exposure. A) Quantitative analysis of cleaved caspase-3-positive apoptotic hepatocytes. Histopathological analysis indicates elevated B) centrilobular hepatocellular necrosis, C) hepatocellular degeneration, and D) inflammatory cell infiltration in all genotypes. Bars indicate the median of scores from individual mice. All comparisons were calculated by Mann-Whitney *U*-test, \**p* < 0.05, \*\**p* < 0.01, \*\*\**p* < 0.001.

Hepatocellular necrosis and degeneration are well-known for their ability to elicit inflammation (Iyer et al., 2009; Sachet et al., 2017; Westman et al., 2019). Analysis shows that the cumulative inflammation score, representing all zones of the hepatic lobules (centrilobular, midzonal, and portal), was markedly enhanced in all mice treated with NDMA (Figure 3D). (Inflammation scores for centrilobular and midzonal regions are shown in Figure S6M and N). Livers from NDMA-treated *Aag*^−/−^ and *AagTg* mice had significantly higher inflammation scores than those of WT animals (Figure 3D). These results suggest that inflammatory cell recruitment is likely mediated by mechanisms dependent on NDMA hepatotoxicity and AAG function. In addition, the significant increase in cytotoxicity and inflammation in *Aag*^−/−^ livers compared to WT supports the hypothesis that unrepaired blocking lesions can produce an overwhelming DNA damage response, leading to toxicity that drives inflammation. It is noteworthy, however, that *Aag*^−/−^ hepatocytes are less susceptible to cytotoxicity, inflammation, and induction of super-bright pan-nuclear γH2AX and MN than *AagTg*, suggesting that blocking lesions are less toxic than SSBs.

### *AagTg* mice are susceptible to NDMA-induced lethality

In addition to acute hepatotoxic lesions, we also observed diminished survival in NDMA-treated *AagTg* mice. Over 12% of *AagTg* mice died within two weeks of injection, whereas nearly all WT and *Aag*^−/−^ mice survived exposure to NDMA (>98%; Table 1). Previous work has shown that acute toxic effects of NDMA occur predominantly in the liver, with minimal evidence of toxicity in other organs in the first few days after exposure (Barnes and Magee, 1954). Thus, elevated animal lethality supports a model wherein *AagTg* livers are particularly sensitized to NDMA toxicity. Our observation that imbalanced BER is lethal is supported by previous studies showing that *AagTg* mice have a significantly lower LD_50_ for alkylating agents compared to WT (Calvo et al., 2016) and *Aag*^−/−^ mice (Calvo et al., 2013).

**Table 1.** *AagTg* mice suffer acute lethality from NDMA exposure.

|  | Total animals<br>injected w/<br>NDMA | Total dead<br>within two<br>weeks | Premature<br>lethality<br>(% injected) |
| --- | --- | --- | --- |
| WT | 137 | 1 | 0.7 |
| *Aag*<sup>-/-</sup> | 93 | 1 | 1.1 |
| *AagTg* | 102 | 13 | 12.7 |

### AAG activity reduces susceptibility to liver cancer

To learn about the impact of AAG levels on cancer risk, we let mice age for ∼10 months to allow time for liver tumors to develop (the average age at necropsy of each saline treatment group was between 10.4–10.6 months, and the average age of each NDMA treatment group was 10.0–10.2 months). We observed a high incidence of macroscopic tumor induction in WT mice, with 67% of NDMA-treated animals developing visibly evident tumors (Table 2). Tumor multiplicity was generally low, wherein no mouse developed more than 5 visible tumors and the median number of tumors was 1 (Figure 4 and Table 2).

**Table 2.** AAG activity modulates liver cancer susceptibility.

|  | Median #<br>tumors/mouse | Incidence (% mice<br>with tumors) |
| --- | --- | --- |
| WT | 1 | 16/24 (67%) |
| *Aag*<sup>-/-</sup> | 4.5 | 19/22 (86%) |
| *AagTg* | 0 | 10/24 (42%) |

**Figure 4.**
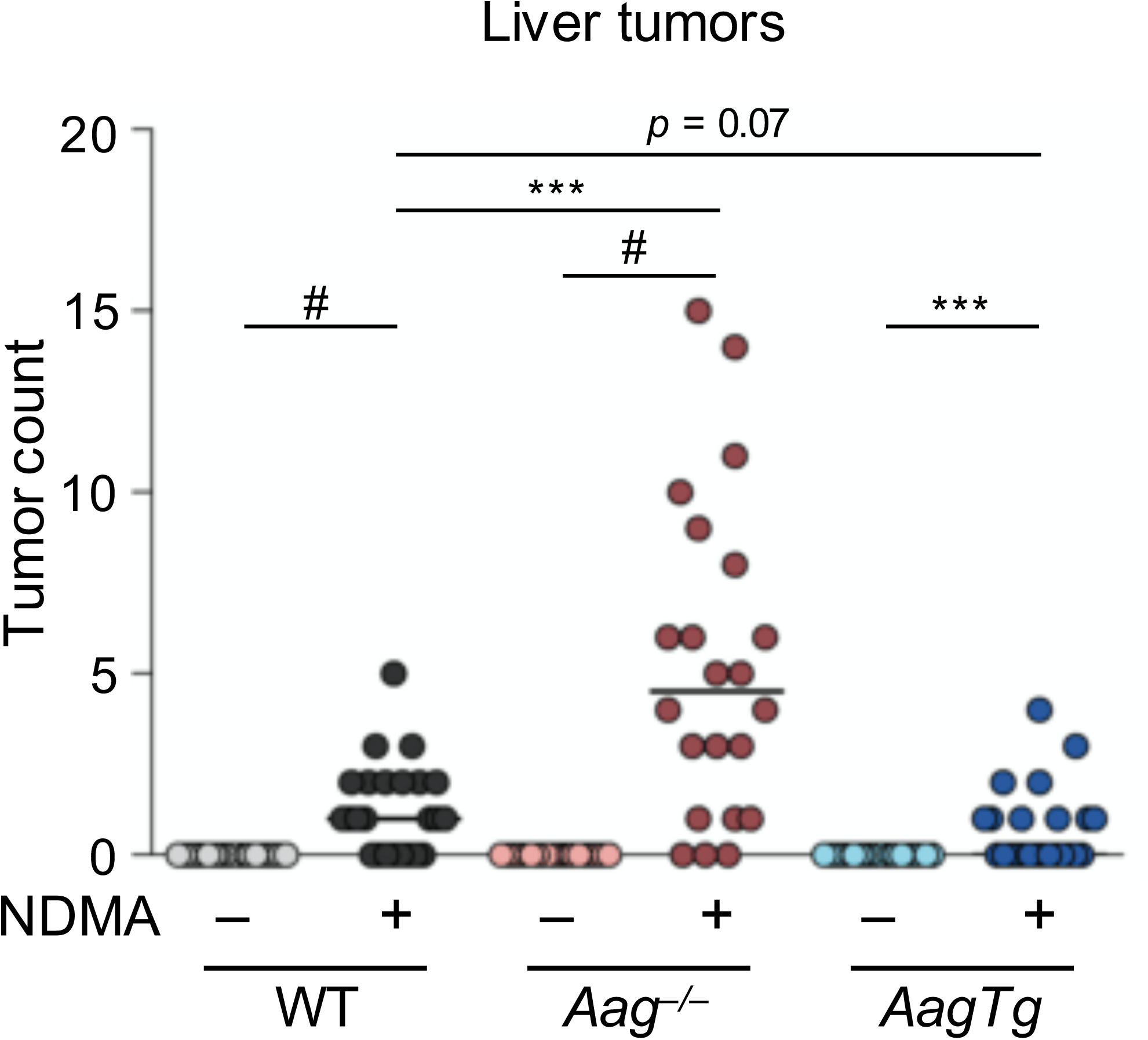
*Aag*^*-/-*^ mice have increased susceptibility to NDMA-induced cancer and *AagTg* mice are resistant. NDMA induced liver cancer in WT, *Aag*^−/−^, and *AagTg* mice. Mice unable to excise the 3MeA lesion (*Aag*^−/−^) are more prone to developing tumors in greater numbers than WT and *AagTg* mice. Mann-Whitney *U*-test, \*\*\**p* < 0.001, ^#^*p* < 0.0001.

*Aag*^−/−^ mice developed significantly more grossly visible tumors than WT mice. In total, 86% of *Aag*^−/−^ mice developed tumors, and the median number of tumors per mouse was significantly higher (4.5 tumors/mouse; Table 2). Considering these data in the context of early phenotypes, it becomes apparent that 3MeA is highly mutagenic and only moderately cytotoxic, which is consistent with survival of cells that acquire tumorigenic mutations. In striking contrast, we observed reduced tumor induction in *AagTg* mice compared to WT (and far lower tumor induction compared to *Aag*^−/−^; Table 2 and Figure 4). Less than half of *AagTg* mice developed any macroscopic tumors, and therefore the median number of tumors was 0 (Table 2). Of note, diminished induction of mutant cells in *AagTg* mice correlated with reduced susceptibility to tumorigenesis.

### Integration of discrete endpoints reveals biomarkers upstream of cancer and lethality

To gain an integrated understanding of how different phenotypes are related to one another, we combined data sets, as shown in Figure 5. Starting from the top and working clockwise, we have plotted key observations from our experiments in chronological order. These include DNA damage (γH2AX foci), replication stress (super-bright pan-nuclear γH2AX), chromosomal instability (micronuclei), cytotoxicity (apoptosis), animal lethality, point mutations, recombination events, and tumor induction. Medians or means were normalized to the highest scoring group for that endpoint (*e*.*g*., NDMA-treated *Aag*^−/−^ mice developed the most tumors [median = 4.5], so all tumor burden medians were normalized to *Aag*^−/−^ NDMA). Light-colored lines show the results for saline-treated mice, and dark colors show the results for NDMA-treated groups.

**Figure 5.**
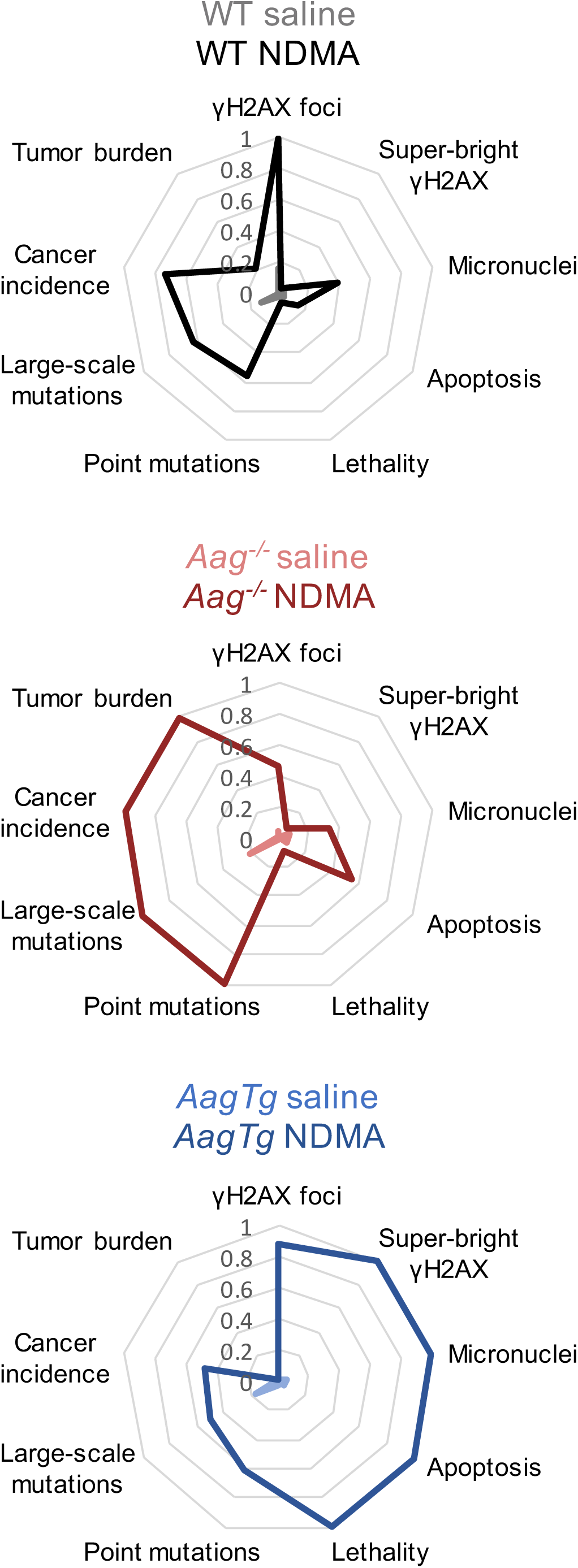
Integration of phenotypic endpoints over time shows that reduced toxicity and increased mutations lie upstream of cancer in *Aag*^−/−^ mice, and DNA damage and cytotoxicity precede lethality in *AagTg* mice. Radar plots showing key endpoints from this study in chronological order. Medians or means from each group for each endpoint are normalized to the highest scoring group at that endpoint (*e*.*g*., NDMA-treated *Aag*^−/−^ mice have the highest tumor burden, so medians of all groups were normalized to the median *Aag*^−/−^. Similarly, NDMA-treated *AagTg* mice showed the highest micronucleus induction, so means of MN results were normalized to the *AagTg* mean.)

Although markers of genotoxicity and animal lethality in NDMA-treated *Aag*^−/−^ mice were lower than or equivalent to those of WT mice, cytotoxicity was significantly increased, and there was very strong induction of mutations and tumors. In stark contrast, overexpression of *Aag* led to poor outcomes shortly after NDMA exposure (including genotoxicity, cytotoxicity, and lethality), and mutations and cancer are low. It is remarkable that for WT mice, the integrated toxicity-related phenotypes (on the right side) as well as the integrated mutagenicity/cancer phenotypes (on the left side) are both reduced compared to *Aag*^−/−^ and *AagTg* mice. In fact, response patterns in WT mice show some similarities to both the *Aag*^−/−^ and the *AagTg* mice, which is consistent with NDMA exposure both saturating AAG excision capacity (leading to persistent 3MeA) while simultaneously causing stress to the downstream BER pathway (leading to increased SSBs and DSBs). Ultimately, the ability to both excise replication blocking lesions and complete the BER pathway mitigates the extreme adverse effects of both replication blocks and strand breaks.

## DISCUSSION

In this work, we have explored the relationships of two fundamentally different classes DNA damage with cancer and lethality, namely lesions that block replication and strand breaks. By varying *Aag* expression, we were able to specifically probe the consequences of unrepaired replication-blocking 3MeA and SSBs formed as BER-intermediates. With a focus on NDMA, a carcinogen that contaminates water, food, and drugs, we found that AAG has an enormous impact on whether cells will survive DNA methylation damage and whether they will eventually develop mutations and cancer. Given that individuals are known to vary in AAG activity, this study points to AAG as being a key variable that dictates the health impact of DNA methylation damage. Methylating agents are not only important environmental contaminants, but they are also used as chemotherapeutic agents to treat cancer. As such, the results of this work have broad relevance to public health as well as to personalized medicine for cancer patients.

We aimed to integrate across multiple phenotypes over time in order to uncover biomarkers with predictive potential for downstream cancer. By presenting the integration of phenotypes in the radar plots, a holistic perspective on biological responses in WT, *Aag*^−/−^ and *AagTg* mice is gained. We found that phenotypes related to toxicity are high in the *AagTg* mice, while downstream mutations and cancer endpoints are equivalent to or lower than WT. In contrast, the *Aag*^−/−^ mice have results heavily skewed toward mutations and cancer with weaker indicators of genotoxicity. These results support a model wherein blocking lesions repaired by AAG are moderately cytotoxic but highly mutagenic, whereas BER intermediates (*e*.*g*., SSBs) are highly toxic, reducing survival of damaged cells and suppressing the development of mutations, but at the cost of increased lethality. Interestingly, strand break-induced toxicity appears to be protective against cancer, consistent with a broad literature showing that disabling apoptosis signaling accelerates cancer (Foster et al., 2012; Lowe and Lin, 2000; Norbury and Zhivotovsky, 2004; Roos and Kaina, 2013). Notably, WT mice show responses that are consistent with both unrepaired 3MeA and SSBs, but each to a lesser extent. Thus, the ability to initiate repair is key to cancer prevention, and completion of the BER pathway is key to prevention of toxicity.

The results of these experiments are consistent with previous studies showing reduced toxicity from 3MeA relative to BER intermediates (Allocca et al., 2019; Calvo et al., 2013; Ebrahimkhani et al., 2014; Kisby et al., 2009; Margulies et al., 2017; Meira et al., 2009; Sobol et al., 2003). In fact, knockout of *Aag* has been shown to protect against alkylation-induced toxicity and degeneration in the retina and cerebellum (Allocca et al., 2019; Calvo et al., 2013; Margulies et al., 2017; Meira et al., 2009), and overexpression of *Aag* sensitizes cells to alkylation-induced toxicity (Hendricks et al., 2002; Ibeanu et al., 1992). In terms of cancer, *Aag*^−/−^ mice have been found to be more susceptible to exposure-induced mutations and tumors in the colon (Calvo et al., 2012; Fahrer et al., 2015; Meira et al., 2008; Wirtz et al., 2010). Interestingly, however, the impact of AAG on mutations appears to depend on context. For example, both overexpression and knockout of *Aag* have been associated with chromosomal aberrations in mammalian cell culture (Coquerelle et al., 1995; Engelward et al., 1998; Engelward et al., 1996; Ensminger et al., 2014; Ibeanu et al., 1992; Kaina et al., 1993). Furthermore, treatment of *Aag*^−/−^ mice with the alkylating agent methyl nitrosourea (which causes the same lesions as NDMA) did not significantly induce sequence rearrangements in the pancreas (Kiraly et al., 2014), suggesting tissue-specific differences in mutagenesis (a phenomenon that has been demonstrated previously (Loktionov et al., 1990; Mientjes et al., 1998; Schmezer et al., 1994; Wang et al., 1998)).

An unexpected pattern revealed by our analyses was the lack of consistency between γH2AX staining and recombination events, despite the fact that DSBs are repaired by HR during S/G2. Interestingly, for *AagTg* mice (wherein increased SSB-induced replication fork breakdown can be restored by mutagenic HR; Figure S7A), we observed similar induction of γH2AX foci as WT, leading one to predict similar degrees of HR induction in WT and *AagTg* mice. However, *AagTg* mice showed a significantly *reduced* frequency of cells harboring HR-driven mutations compared to WT. Analyses of super-bright pan-γH2AX staining (indicative of toxic replication stress), MN induction, apoptosis, and necrosis all suggest that damaged cells are being eliminated rather than surviving with mutations. On the other hand, livers from *Aag*^−/−^ mice showed significantly reduced γH2AX staining compared to WT, and yet they developed significantly more HR-driven mutations. The susceptibility of *Aag*^−/−^ mice to mutagenic recombination may therefore reflect DSB-independent HR events. While several studies have linked replication blocking lesions to HR through an endonuclease that creates DSBs (Hanada et al., 2007; Hanada et al., 2006; Saugar et al., 2013; Willis and Rhind, 2009), direct evidence of blocking lesions leading to fork breakdown and free DSB ends is lacking, in part because the predominant method for studying so-called blocked replication forks is to deplete cells of nucleotides, which is quite different from inducing damage that inhibits polymerases. Indeed, blocking lesions are known to induce template switching (Ait Saada et al., 2018; Lambert et al., 2010; Marians, 2018), and our data suggest that this is likely the dominant response to blocking lesions. In the case of the direct repeat HR substrate used for these studies, fork reversal (caused by leading strand blockage) may lead to misalignments that reconstitute the full-length EGFP cDNA (Figure S7B). Alternatively, lagging strand blockage can be bypassed by invasion of the sister chromatid, and misalignment during this process can also restore full-length EGFP (Figure S7C).

An exciting and novel aspect of this study is that the combination of *gpt* delta and RaDR transgenes enables analyses of both point mutations and mutagenic recombination events within the same tissues. RaDR mutations can be evaluated by fluorescent microscopy on fresh tissue in mere minutes, and following imaging, the tissues can be flash frozen for subsequent *gpt* point mutation analyses. Using our dual detection approach, we show that conditions that promote point mutations also promote sequence rearrangements, an observation that has been suggested by many diverse studies but has proven difficult to demonstrate directly. Importantly, the results of this study show strong predictive capacity of RaDR and *gpt* mutation assays for cancer risk, suggesting these assays may serve as effective surrogates for time- and resource-intensive carcinogenicity studies, especially for DNA damaging agents. Furthermore, results indicate that genotoxicity analyses, such as γH2AX and MN, are not as effective in predicting cancer risk as analyses of mutations, and it may even be misleading if not combined with analyses of toxicity.

A major challenge in chemotherapy is the risk of mutations that drive secondary cancers (Choi 2014). Ideally, a drug or combination of drugs would be toxic without being mutagenic. The approaches described here present an interesting opportunity to optimize cancer chemotherapeutics by integrating multiple early stage biomarkers predictive of both toxicity and downstream cancer. Evaluation of mutagenicity with RaDR;*gpt* mice in combination with analyses of DNA damage (*e*.*g*., γH2AX and MN) and toxicity may prove to be an effective strategy for optimizing treatments so as to avoid secondary cancers before going to clinical trials. This is particularly important for childhood cancer therapies, for which secondary cancers can arise over a decade after treatment.

In addition to replication fork breakdown-associated DSBs that signal apoptosis, prior studies show that overactivity of PARP is a significant modulator of AAG-driven toxicity due to increased strand breaks and PARP-1 hyperactivation (Alhumaydhi et al., 2020; Allocca et al., 2017; Allocca et al., 2019). PARP-1 stimulates repair of SSBs by polymerizing ADP-ribose from NAD+, so an excess of DNA damage causes NAD+ depletion and a form of programmed necrotic cell death termed parthanatos (Xu et al., 2006; Yu et al., 2002; Zhao et al., 2018). We have previously shown that *AagTg* mice experience elevated cytotoxicity and tissue degeneration from DNA alkylating treatments in the cerebellum, retina, spleen, thymus, bone marrow, and pancreas (Allocca et al., 2017; Allocca et al., 2019; Calvo et al., 2013; Margulies et al., 2017), and that this is often dependent on PARP-1. Indeed, histopathology of NDMA-treated *AagTg* livers showed significantly elevated necrosis over that of WT and *Aag*^−/−^ livers. It is also noteworthy that PARylation by PARP-1 facilitates the recruitment of several proteins involved in HR (Bryant et al., 2009; Haince et al., 2008; Li and Yu, 2013); as a result, NAD+ depletion in *AagTg* cells may have prevented effective HR, leading to the observed reduction in HR-driven mutations in those mice. Since high levels of cytotoxicity drive degenerative diseases and aging, individuals with high AAG activity may be at elevated risk for such pathologies. In this context, it is interesting to note that *AagTg* mice have approximately 6-fold higher AAG activity than WT in the liver (Calvo et al., 2013), whereas AAG activity can vary by as much as 20-fold among people (Calvo et al., 2013; Crosbie et al., 2012; Hall et al., 1993), calling attention to the biological relevance of this study.

Importantly, the analyses presented here grouped equal numbers of males and females of the same genotype. However, it is known that males and females differ in susceptibility to alkylation damage (Allocca et al., 2017; Allocca et al., 2019; Likhite et al., 2004) and liver cancer (Hassan et al., 2017; Naugler et al., 2007; Rao and Vesselinovitch, 1973). A brief report on sex differences observed in these experiments is in preparation, but further studies of sex-dependent differences in *Aag*^−/−^ and *AagTg* mice will help define interindividual variability in responses to NDMA-induced health outcomes.

The work presented here uncovers a variety of insights regarding human susceptibility to disease. NDMA contamination is widespread, and there are many sources of human exposure. Since people have been shown to vary substantially in AAG activity (Calvo et al., 2013; Chaim et al., 2017; Crosbie et al., 2012; Leitner-Dagan et al., 2012), our research may provide a means of stratifying exposed populations into low- and high-risk categories for different adverse health outcomes. For example, DNA methylating agents are often used for cancer chemotherapy, and so knowledge of the levels of AAG in a tumor can be used for precision medicine, since high levels of AAG would predict effective cytotoxicity, whereas low AAG would point to increased mutations (and potentially more aggressive tumors). Importantly, from the public health perspective, novel technologies that enable screening of people for their AAG activity (Chaim et al., 2017; Nagel et al., 2014) are paving the way for precision prevention. Taken together, the results of this study provide a basis for advancements in predicting outcomes of exposure to DNA alkylating agents that promise to help in both treating and preventing cancer.

## Acknowledgements

This work was supported by the National Institute of Environmental Health Sciences (NIEHS) Superfund Basic Research Program, National Institutes of Health (NIH), P42-ES027707; NIEHS Core Center Grant, NIH, P30-ES002109; NIEHS Toxicology Training Grant, NIH T32-ES007020; NIH Grant R01-CA080024; NIEHS Small Business Innovation Research Grant, NIH R44-ES0264644; and NIH Biomedical Technology Research Resource Grant 5-P41EB015871-32, and the Center for Advanced Imaging at Harvard University.

We thank Caroline Atkinson and Joanna Richards in the MIT Division of Comparative Medicine for H&E and cleaved caspase-3 staining and Kathleen Cormier in the MIT Koch Institute Histology Core for Aperio slide scanning of caspase-stained slides. We appreciate the assistance of Aimee Moise in necropsies and RaDR imaging and Judy Yau in *gpt* analysis. We thank Jeff Dixon for creating the Template Switching and Fork Breakdown diagrams.

## Author Contributions

Conceptualization, JEK, FW, LDS, JME, and BPE; Methodology, JEK, JJC, DNW, BPE, FW, LDS, JME, SDD; Resources, RGC, BPE, LDS, JME; Software, DNW; Investigation, JEK, JJC, ALA, INK, SEC, ISN, DKT, SLA; Data curation, JEK and JJC; Writing – Original Draft, JEK and BPE; Writing – Review & Editing, JEK, SEC, ALA, JJC, INK, SDD, FMW, LDS, BPE; Visualization, JEK, JJC, IK, ALA, SEC; Supervision, BPE, JME, FMW, LDS, SDD; Funding Acquisition, BPE, JME, LDS, FMW, SDD, DNW.

**STAR Methods** (https://www.cell.com/star-authors-guide)

## RESOURCE AVAILABILITY

*Lead Contact (*responsible for communication with the journal before and after publication and is the arbiter of disputes, including concerns related to reagents or resource sharing. Authors must be willing to distribute all materials, datasets, and protocols used in the manuscript.)

Further information and requests for resources and reagents should be directed to and will be fulfilled by the Lead Contact, Bevin Engelward.

### Materials Availability

This study did not generate new unique reagents.

### Data and Code Availability

The mass spectrometry proteomics data are available at the ProteomeXchange Consortium via the PRIDE (Perez-Riverol et al., 2019) partner repository with the dataset identifier PXD021142.

## EXPERIMENTAL MODEL AND SUBJECT DETAILS

### Animals

All experimental mice were on a C57BL6 genetic background. *RaDR*^*R/R*^;*gpt*^*g/g*^ mice were created by crossing RaDR-GFP mice (B6.129S4(Cg)-*Gt(ROSA)26Sor*^*tm1(CAG-EGFP*)Bpeng*^) (Sukup-Jackson et al., 2014) with *gpt* delta mice (Nohmi et al., 1996) (a gift from T. Nohmi). *RaDR*^*R/R*^;*gpt*^*g/g*^;*Aag*^*-/-*^ mice were generated by crossing *RaDR*^*R/R*^;*gpt*^*g/g*^ and *Aag*^*-/-*^ mice (described previously (Engelward et al., 1997)). *RaDR*^*R/R*^;*gpt*^*g/g*^;*AagTg* were generated by crossing *RaDR*^*R/R*^;*gpt*^*g/g*^ and *AagTg* mice (described previously (Calvo et al., 2013)). The mice were maintained in AAALAC-certified animal care facilities and provided standard food and water *ad libitum*. All animal procedures were performed according to the NIH Guide for the Care and Use of Laboratory Animals and protocols approved by the Massachusetts Institute of Technology Committee on Animal Care.

## METHOD DETAILS

### NDMA synthesis

NDMA was prepared as previously described (He et al., 2019; Heath, 1961). Briefly, sodium nitrite in water was slowly added to a solution of dimethylanime in water and acetic acid. After cooling, 10N NaOH was added and the solution was extracted four times with ether. The ether solution was dried over Na_2_SO_4_ and NDMA isolated by fractional distillation having a boiling point of 148-149°C. The pale yellow liquid was characterized by NMR and mass spectrometry: MS, ESI m/z, 75.0056 (M+H); ^1^H 300 MHz NMR, CDCl_3_ δ 3.06 (s, 3H,Me *cis*), 3.76 (s,3H,Me *trans*). The NMR assignments are in agreement with those reported for NDMA by Karabatsos and Taller (Karabatsos, 1964). Aliquots of NDMA were packaged in sealed ampules under Ar gas and stored at −20°C.

### NDMA treatment

Litters of mice were designated for saline or NDMA treatment at birth. A total dose of 10.5 mg/kg NDMA diluted in saline was administered intraperitoneally over two separate injections, according to (Dass et al., 1998). One-third of the dose (3.5 mg/kg NDMA in saline, 10 μL volume) was given at 8 days of age, and the remaining two-thirds of the dose (7 mg/kg, 20 μL volume) was given at 15 days of age. Control mice were sham treated at the same timepoints with equivalent volumes of saline.

Mice were euthanized by asphyxiation with carbon dioxide according to AVMA guidelines and necropsied 24 hours (phosphoproteomics, immunostaining, and histopathology), 48 hours (flow cytometric micronucleus and proliferation), 10 weeks (mutations and histopathology), or 9 to 11 months (tumors) post-second injection. Whenever possible, equal numbers of males and females of each genotype and treatment group were analyzed for each endpoint. When data were collected from an excess number of males or females for an endpoint, individual data points were selected by random number generator for exclusion from analysis.

### Gpt assay

Samples of liver tissue were collected after RaDR imaging, flash frozen in liquid nitrogen, and stored at −80°C until analysis (n = 5 males and 5 females from each group). Mutations in the *gpt* gene were identified by selection with 6-thioguanine (6-TG), as previously described (Chawanthayatham et al., 2017; Nohmi et al., 1996). Briefly, liver tissues were pulverized with a mortar and pestle with liquid nitrogen. Genomic DNA was extracted from approximately 25 mg of liver tissue using the RecoverEase DNA Isolation Kit (Agilent Technologies). The λ-EG10 phage were packaged *in vitro* from genomic DNA using Transpack packaging extract (Agilent Technologies). The λ-EG10 phage were then transfected into *Escherichia coli* YG6020 expressing Cre-recombinase, generating a 6.4-kb plasmid carrying the *gpt* and chloramphenicol acetyltransferase genes. These bacteria were cultured on selective media containing chloramphenicol and 6-TG or chloramphenicol alone. 6-TG resistance was confirmed by regrowth of colonies on plates containing chloramphenicol and 6-TG. The samples were processed and analyzed in a blinded fashion.

### Phosphoproteomic analysis

Twenty-four hours after the second injection, animals were humanely euthanized and portions of livers were immediately excised, flash frozen in liquid nitrogen, and stored at −80°C until analysis (WT mice only; saline, one male and one female; NDMA, three females). Liver tissues were homogenized in ice-cold 8M Urea (Sigma) with three 10-second pulses. Proteins were processed, digested and desalted as described previously (Dittmann et al., 2019). Lyophilized peptide aliquots of 400 µg (of starting protein) were labeled with TMT10-plex labeling kits (Thermo Fisher). Phosphopeptides were enriched with a 2-step enrichment process consisting of immunoprecipitation (IP) followed by Fe-NTA-based immobilized metal affinity chromatography (IMAC). TMT-labeled samples were resuspended in IP buffer (100 mM Tris-HCl, 1% Nonidet P-40, pH 7.4) and incubated overnight with PTMScan Phospho-ATM/ATR Substrate Motif kit (Cell Signaling Technology). Peptides were eluted twice, each with 25 µL of 0.2% trifluoroacetic acid (TFA) for 10 minutes at room temperature followed by a secondary Fe-NTA-based IMAC to remove non-specifically retained non-phosphopeptides. High-Select Fe-NTA enrichment kit (Pierce) was used according to manufacturer’s protocol with following modifications. After washing the Fe-NTA spin columns, beads were resuspended in 25 µL of binding washing buffer. Eluates from IP were incubated with Fe-NTA beads for 30 minutes. Peptides were eluted twice with 20 µL of elution buffer into a 1.7-mL microcentrifuge tube. Eluates were dried in SpeedVac until 1-5 µL of sample remained. Samples were resuspended in 10 µL of 5% acetonitrile in 0.1% formic acid and loaded directly onto an in-house packed analytical capillary column (50 µm ID x 10 cm) packed with 5-µm C18 beads (YMC gel, ODS-AQ, AQ12S05).

Liquid chromatography tandem mass spectrometry (LC-MS/MS) of phosphopeptides and crude lysate analysis was carried out on an Agilent 1260 LC coupled to a Q Exactive HF-X mass spectrometer (Thermo Fisher) as described previously (Dittmann et al., 2019). Raw mass spectra data files were processed with Proteome Discoverer version 2.2 (Thermo Fisher) and searched against the mouse SwissProt database using Mascot version 2.4 (Matrix Science). TMT reporter quantification was extracted using Proteome Discoverer. MS/MS spectra were searched with the following settings: mass tolerance of 10 ppm for precursor ions; 20 mmu for fragment ions; fixed modification for cysteine carbamidomethylation, TMT-labeled lysine, TMT-labeled peptide N-termini; dynamic modifications for methionine oxidation and phosphorylation of serine, threonine and tyrosine. Peptide spectrum matches (PSMs) were filtered according to following parameters: rank = 1, search engine rank = 1, mascot ion score > 20, isolation interference < 30%, average TMT signal > 1000. Peptides with missing values across any channel were filtered out. Phosphorylation sites were localized using ptmRS module (Taus et al., 2011) on Proteome Discoverer. PSMs with >95% localization probability were included for further analysis. Only peptides containing ‘SQ’ or ‘TQ’ sequence motif were included for final analysis. Peptide quantification was normalized with relative median values obtained from crude peptide analysis. Further data analysis was performed in Python (version 3.6) and MATLAB (R2016a).

### RaDR analysis

Livers were collected from mice 10 weeks after the second NDMA injection (WT saline n = 12 males, 12 females; WT NDMA n = 11 males, 11 females; *Aag*^−/−^ saline n = 11 males, 11 females; *Aag*^−/−^ NDMA n = 12 males, 12 females; *AagTg* saline n = 13 males, 13 females; *AagTg* NDMA = 9 males, 9 females). Freshly excised livers were held on ice in 0.01% trypsin inhibitor (Boston BioProducts) in PBS prior to imaging. The entire left lobe of the liver was secured between a glass slide and a cover slip. The dorsal surface of each liver was then imaged with a Nikon Eclipse Ti2 scanning microscope on the 2x objective in the FITC channel using an Andor Zyla 4.2 camera and NIS Elements software. A user-trained two-stage machine learning algorithm was used to identify and enumerate fluorescent foci within intact tissue. Similar to our previous work (Wadduwage et al., 2018), we first identified potential regions (termed region proposals) that might contain foci, and then classified them into True Foci or False Regions (Figure S4A). Two deep convolutional neural networks (DCNNs) were used that were trained using 10 manually annotated images for both tasks. The first network (i.e. Region Proposal Network), segmented the foci-like regions. Then we used their locations to extract the corresponding image patches (i.e. region proposals). These region proposals were then fed to a second DCNN (i.e. Proposal Classifier Network) that classified them into either True Foci, or False Regions. The true foci locations were then listed and counted to get the final foci count. The 10 manually annotated training images comprised samples from each genotype and treatment group. All liver images were analyzed by the machine learning program based on parameters developed from training data. The number of fluorescent foci was normalized to the area of the liver in the image.

### Histological analysis

Sections of liver were fixed in 10% buffered formalin, embedded in paraffin, and sectioned at 4-μm thickness using a microtome, followed by hematoxylin and eosin (H&E) staining. Liver sections from 4 males and 4 females of each group (24h and 10-week timepoints) were scored by a board-certified veterinary pathologist blinded to sample identity. Specific lesions were graded with a numerical score from 0 to 4, in which 0 = normal, 1 = minimal, 2 = mild, 3 = moderate, and 4 = severe. The following hepatic lesions were graded: inflammation, hepatocellular degeneration, hepatocellular necrosis, nuclear enlargement (karyomegaly), foci of hepatocellular alteration, hepatic lipidosis, extramedullary hematopoiesis, Kupffer cell hyperplasia, Ito cell hyperplasia, bile duct hyperplasia/dysplasia, and fibrosis (Thoolen et al., 2010). Lesions for the liver scored as present (1) or absent (0) included hemorrhage and neoplasia. A total inflammation score for each liver was generated by combining individual scores of portal, midzonal, and centrilobular inflammation from each submitted section (Snider et al., 2018). Foci of altered hepatocytes were classified based on morphologic criteria reported by Thoolen and colleagues (Thoolen et al., 2010). Sections were examined using an Olympus BX41 microscope attached with an iKona digital camera and photographed.

### γ*H2AX staining and analysis*

Formalin-fixed tissue sections (4 μm) from 4 males and 4 females from each treatment group (24h timepoint) were deparaffinized in xylenes, rehydrated, and subjected to heat-induced epitope retrieval (HIER) in citrate buffer pH 6.1 (Agilent). Sections were blocked with 5% bovine serum albumin (BSA) and 0.5% Tween 20 for 1 hour at room temperature, then incubated with an antibody for γH2AX (1:200; Cell Signaling Technologies) in 1% BSA and 0.5% Tween 20 in PBS overnight at 4°C. Sections were then washed and incubated with a secondary antibody conjugated to an AlexaFluor 488 probe (1:400; Invitrogen) for 1 hour at room temperature. Nuclei were counterstained with DAPI with ProLong Gold AntiFade (ThermoFisher). Stained tissue sections were imaged at 60x under DAPI and FITC filters using a Nikon Eclipse Ti2 microscope and an Andor Zyla 4.2 camera. A complete immunofluorescence staining protocol is available at https://nextgen-protocols.org/protocol/staining-for-%ce%b3h2ax-in-paraffin-embedded-tissue-sections/.

Cells containing 5 or more γH2AX foci or pan-nuclear staining were manually annotated and enumerated in a blinded fashion. Super-bright pan-nuclear staining was determined with ImageJ (NIH) by setting a conservative pixel intensity threshold and excluding particles smaller than 10 μm^2^. Nuclei were identified by using Ilastik (Berg et al., 2019) and KNIME (University of Konstanz, Zurich, Switzerland) software to segment nuclei based on user-annotated training images. Segmented particles were counted in ImageJ for an approximate number of nuclei in the image, and the percentage of nuclei falling into each category (foci, pan-nuclear, super-bright pan-nuclear) was calculated. Data were collected for at least 1,300 nuclei from each mouse.

### Caspase staining and analysis

Formalin-fixed tissue sections (4 μm) from 4 males and 4 females from each treatment group (24h timepoint, same mice as those analyzed for γH2AX and histopathology) were deparaffinized in xylenes, rehydrated, and subjected to HIER in Buffer H, pH 8.8 (Thermo Scientific). After cooling, endogenous peroxidase was blocked with Peroxidazed 1 (Biocare Medical) for 5 minutes and slides were blocked in Background Sniper (Biocare Medical) for 15 minutes. Slides were then incubated with cleaved caspase-3 primary antibody (Cell Signaling Technology) for 60 minutes at room temperature, washed, and incubated with Rabbit-on-Rodent HRP Polymer (Biocare Medical) for 30 minutes at room temperature. Slides were washed and incubated with DAB (Betazoid DAB Chromagen kit, Biocare Medical), washed, and counterstained with hematoxylin. Stained slides were scanned with a Leica Aperio AT2 slide scanning microscope and analyzed with QuPath software (Bankhead et al., 2017). After blinding filenames, 4 equal-sized, randomly selected, independent regions of tissue were annotated for number of nuclei and number of apoptotic events. The percentage of apoptotic events for each animal was calculated from the total number of apoptotic hepatocytes and total number of nuclei from four representative 400x fields.

### Micronucleus assay

Supplies for collecting mouse livers at MIT and shipping to Litron for flow cytometric hepatocyte micronucleus (MNHEP) scoring were from Prototype In Vivo MicroFlow^®^ BASIC ML Kits, Litron Laboratories and included Liver Preservation Buffer Solution, 2 ml vials and ExaktPak shipping containers with ice packs. Reagents and supplies used at Litron for preparing livers for MNHEP scoring were from Prototype In Vivo MicroFlow^®^ PLUS ML Kits, Litron Laboratories and included Liver Rinse, EGTA Solution, Collagenase Solution, Incomplete Lysis Solution 1, Incomplete Lysis Solution 2, anti-Ki67-eFluor® 660, Liver Nucleic Acid Dye (contains SYTOX^®^ Green), and RNase Solution.

Mice were euthanized 48 hours after the second NDMA injection and livers were collected into 1 mL ice-cold Liver Preservation Buffer, packed on ice, and shipped overnight to Litron Laboratories. Upon receipt at Litron, each liver was removed from the Liver Preservation Buffer, patted dry, placed into a separate flask containing 10 mL of Liver Rinse Solution, and processed as described previously (Avlasevich et al., 2018). Samples were analyzed with a FACSCanto™ II flow cytometer equipped with 488 and 633 nM excitation (BD Biosciences, San Jose, CA).

Instrumentation settings and data acquisition/analysis were controlled with FACSDiva™ software v6.1.3 (BD Biosciences). SYTOX Green-associated fluorescence emissions were collected in the FITC channel (530/30 band-pass filter), and anti-Ki67-eFluo^R^ 660-associated fluorescence emissions were collected in the APC channel (660/20 band-pass filter). The flow cytometry gating strategy for MNHEP scoring required events to fall within each of three regions and one histogram marker before they were scored as nuclei or micronuclei. The incidence of flow cytometry-scored MNHEP is expressed as frequency percent (no. micronuclei/no. nuclei x 100) of 20,000 SYTOX Green positive nuclei per specimen. Simultaneous with micronucleus assessments, an experimental index of hepatocyte proliferation was collected based on gating for Ki67-positive nuclei.

MNHEP microscopy was performed by adding SYTOX Green-stained cells to acridine orange-coated slides, then imaged on an Olympus BH-2 microscope with a 40x objective as previously described (Avlasevich et al., 2018).

### Tumor analysis

Gross surface lesions on the entire liver were recorded at necropsy. Distinct macroscopic tumors were enumerated; multifocal to coalescing tumorous regions were counted as one.

## QUANTIFICATION AND STATISTICAL ANALYSES

Statistical analyses were performed using the GraphPad Prism software. Tumor multiplicity, RaDR foci, *gpt* mutant fractions, caspase staining, and histopathology scores were compared by Mann-Whitney *U*-test. Phosphoproteome, micronucleus, γH2AX, and Ki67 analyses were compared by unpaired Student’s *t*-test. A *P* value was considered significant if less than 0.05.

## Supplemental information titles and legends

**Figure S1. NDMA treatment regimen and collection timepoints**. A total of 10.5 mg/kg NDMA was injected in a split dose in neonatal mice, with 3.5 mg/kg injected at 8 days of age and 10 mg/kg injected at 15 days of age. Control mice were injected with equivalent volumes of saline at the same timepoints. Analyses were performed 24h, 48h, 10 weeks, and 9-11 months after the second injection.

**Figure S2. Livers of NDMA-treated mice show evidence of hepatocellular degeneration and hypertrophy 10 weeks after exposure, but not neoplasia**. A &D) Representative hematoxylin and eosin (H&E) sections of liver from WT mice showed centrilobular (squares) to midzonal areas of hepatocellular degeneration and hypertrophy with and without karyomegaly (arrowheads) and intranuclear invaginations. Similar hepatocellular changes were noted in sections of livers from NDMA-treated *Aag*^−/−^ (F & H) and *AagTg* mice (J & L). Sections of livers from control WT mice (A & C), *Aag*^−/−^ (F & H), and *AagTg* (I & K) were within normal limits. Kupffer cell aggregates were present in control and treated groups (arrows). No significant foci of hepatocellular alteration were observed in any tissue. Original magnification x400, scale bars = 50 μm.

**Figure S3. RaDR mice enable visualization of mutant cells in the liver by fluorescence microscopy**. A) Whole-mount microscopy of EGFP in untreated RaDR null and *RaDR*^R/R^ livers. Spontaneous recombinogenic mutations are visible as fluorescent foci in RaDR tissues, but not in mice without the RaDR transgene. Scale bar = 5 mm. B) Mutant hepatocyes are visible in an overlay of EGFP and H&E. Scale bar = 20 μm. Hepatocyte H&E overlay reused from Sukup-Jackson et al., PLOS Genetics, 2014, https://doi.org/10.1371/journal.pgen.1004299 under Creative Commons license CC BY 4.0. This image was cropped from a larger figure.

**Figure S4. Machine learning network for automated enumeration of fluorescent foci in RaDR tissues**. A) Schematic of foci counting algorithm. See STAR methods for details. B) Sample RaDR liver image (top) and foci counting algorithm output (bottom).

**Figure S5. Proliferation is not significantly different among NDMA-treated mice 48h post-exposure**. The proportion of proliferating cells was determined by flow cytometry of Ki67-positive cells. Data represented as mean ± SD. For all graphs, each data point represents one mouse. Statistical comparisons calculated by unpaired Student’s *t*-test, \**p* < 0.05, \*\**p* < 0.01, ^#^*p* < 0.0001.

**Figure S6. Acute periacinar hepatocellular damage is more pronounced in *Aag*Tg mice at 24h post-NDMA treatment**. A–D) Representative H&E staining of liver sections from WT mice show individual necrotic hepatocytes (arrowheads) and numerous degenerate hepatocytes surrounding a central vein in NDMA treated mice (B&D). E–H) Sections of liver from *Aag*^−/−^ mice exhibited multifocal areas of mild centrilobular necrosis and degeneration (arrows) with low numbers of macrophages and fewer neutrophils at the periphery. I–L) Sections of liver from *AagTg* mice show larger areas of centrilobular (periacinar) hepatocellular necrosis (asterisk) and degeneration surrounded by low numbers of macrophages. No histopathologic changes were noted in sections of liver from control WT (A & C), *Aag*^−/−^ (F & H), and *AagTg* mice (I & K). Original magnification x400, scale bars = 50 μm. M & N) Leukocytic infiltration scores in centrilobular (M) and midzonal (N) regions in the liver. Bars indicate the median of scores from individual mice. Mann-Whitney *U*-test, \**p* < 0.05, \*\**p* < 0.01, \*\*\**p* < 0.001, ^#^*p* < 0.0001.

**Figure S7. Mechanisms to produce EGFP within the RaDR transgene at replication forks**. A) Fork breakdown at an SSB can be restored by HR to reconstitute full-length EGFP cDNA (green segments) within the RaDR transgene (black boxes indicate deleted sequences). B) Replication blocks in the leading strand can lead to template switching by fork reversal, potentially leading to production of full-length EGFP in the RaDR transgene. C) Replication blocks in the lagging strand can be bypassed by utilizing the nascent strand on the sister chromatid as a template for synthesis. Template switching potentially leading to production of full-length EGFP in the RaDR transgene (D-loop in step 2 not shown).

**Table S1. Processed phosphoproteomic data**.

## References

Aissi-Ben Moussa, S., A. Moussa, T. Lovecchio, N. Kourda, T. Najjar, S. Ben Jilani, A. El Gaaied, N. Porchet, M. Manai, and M.P. Buisine. 2009. Identification and characterization of a novel MLH1 genomic rearrangement as the cause of HNPCC in a Tunisian family: evidence for a homologous Alu-mediated recombination. Familial cancer. 8:119–126.

Ait Saada, A., S.A.E. Lambert, and A.M. Carr. 2018. Preserving replication fork integrity and competence via the homologous recombination pathway. DNA repair. 71:135–147.

Alhumaydhi, F.A., O.L.D. de, D.L. Bordin, A.S.M. Aljohani, C.B. Lloyd, M.D. McNicholas, L. Milano, C.F. Charlier, I. Villela, J.A.P. Henriques, K.E. Plant, R.M. Elliott, and L.B. Meira. 2020. Alkyladenine DNA glycosylase deficiency uncouples alkylation-induced strand break generation from PARP-1 activation and glycolysis inhibition. Sci Rep. 10:2209.

Allocca, M., J.J. Corrigan, K.R. Fake, J.A. Calvo, and L.D. Samson. 2017. PARP inhibitors protect against sex- and AAG-dependent alkylation-induced neural degeneration. Oncotarget. 8:68707–68720.

Allocca, M., J.J. Corrigan, A. Mazumder, K.R. Fake, and L.D. Samson. 2019. Inflammation, necrosis, and the kinase RIP3 are key mediators of AAG-dependent alkylation-induced retinal degeneration. Sci Signal. 12.

Anwar, S.L., W. Wulaningsih, and U. Lehmann. 2017. Transposable Elements in Human Cancer: Causes and Consequences of Deregulation. International journal of molecular sciences. 18.

Avlasevich, S.L., S. Khanal, P. Singh, D.K. Torous, J.C. Bemis, and S.D. Dertinger. 2018. Flow cytometric method for scoring rat liver micronuclei with simultaneous assessments of hepatocyte proliferation. Environmental and molecular mutagenesis. 59:176–187.

Bankhead, P., M.B. Loughrey, J.A. Fernandez, Y. Dombrowski, D.G. McArt, P.D. Dunne, S. McQuaid, R.T. Gray, L.J. Murray, H.G. Coleman, J.A. James, M. Salto-Tellez, and P.W. Hamilton. 2017. QuPath: Open source software for digital pathology image analysis. Sci Rep. 7:16878.

Baritaud, M., L. Cabon, L. Delavallee, P. Galan-Malo, M.E. Gilles, M.N. Brunelle-Navas, and S.A. Susin. 2012. AIF-mediated caspase-independent necroptosis requires ATM and DNA-PK-induced histone H2AX Ser139 phosphorylation. Cell death & disease. 3:e390.

Barnes, J.M., and P.N. Magee. 1954. Some toxic properties of dimethylnitrosamine. Br J Ind Med. 11:167–174.

Beckta, J.M., S.M. Dever, N. Gnawali, A. Khalil, A. Sule, S.E. Golding, E. Rosenberg, A. Narayanan, K. Kehn-Hall, B. Xu, L.F. Povirk, and K. Valerie. 2015. Mutation of the BRCA1 SQ-cluster results in aberrant mitosis, reduced homologous recombination, and a compensatory increase in non-homologous end joining. Oncotarget. 6:27674–27687.

Bekker-Jensen, S., and N. Mailand. 2010. Assembly and function of DNA double-strand break repair foci in mammalian cells. DNA repair. 9:1219–1228.

Beranek, D.T. 1990. Distribution of methyl and ethyl adducts following alkylation with monofunctional alkylating agents. Mutation research. 231:11–30.

Berg, S., D. Kutra, T. Kroeger, C.N. Straehle, B.X. Kausler, C. Haubold, M. Schiegg, J. Ales, T. Beier, M. Rudy, K. Eren, J.I. Cervantes, B. Xu, F. Beuttenmueller, A. Wolny, C. Zhang, U. Koethe, F.A. Hamprecht, and A. Kreshuk. 2019. ilastik: interactive machine learning for (bio)image analysis. Nat Methods. 16:1226–1232.

Bishop, A.J., and R.H. Schiestl. 2001. Homologous recombination as a mechanism of carcinogenesis. Biochimica et biophysica acta. 1471:M109–121.

Bryant, H.E., E. Petermann, N. Schultz, A.S. Jemth, O. Loseva, N. Issaeva, F. Johansson, S. Fernandez, P. McGlynn, and T. Helleday. 2009. PARP is activated at stalled forks to mediate Mre11-dependent replication restart and recombination. The EMBO journal. 28:2601–2615.

Calvo, J.A., M. Allocca, K.R. Fake, S. Muthupalani, J.J. Corrigan, R.T. Bronson, and L.D. Samson. 2016. Parp1 protects against Aag-dependent alkylation-induced nephrotoxicity in a sex-dependent manner. Oncotarget. 7:44950–44965.

Calvo, J.A., L.B. Meira, C.Y. Lee, C.A. Moroski-Erkul, N. Abolhassani, K. Taghizadeh, L.W. Eichinger, S. Muthupalani, L.M. Nordstrand, A. Klungland, and L.D. Samson. 2012. DNA repair is indispensable for survival after acute inflammation. The Journal of clinical investigation. 122:2680–2689.

Calvo, J.A., C.A. Moroski-Erkul, A. Lake, L.W. Eichinger, D. Shah, I. Jhun, P. Limsirichai, R.T. Bronson, D.C. Christiani, L.B. Meira, and L.D. Samson. 2013. Aag DNA glycosylase promotes alkylation-induced tissue damage mediated by Parp1. PLoS genetics. 9:e1003413.

Chaim, I.A., Z.D. Nagel, J.J. Jordan, P. Mazzucato, L.P. Ngo, and L.D. Samson. 2017. In vivo measurements of interindividual differences in DNA glycosylases and APE1 activities. Proceedings of the National Academy of Sciences of the United States of America. 114:E10379–E10388.

Chanoux, R.A., B. Yin, K.A. Urtishak, A. Asare, C.H. Bassing, and E.J. Brown. 2009. ATR and H2AX cooperate in maintaining genome stability under replication stress. The Journal of biological chemistry. 284:5994–6003.

Charych, D.H., M. Coyne, A. Yabannavar, J. Narberes, S. Chow, M. Wallroth, C. Shafer, and A.O. Walter. 2008. Inhibition of Cdc7/Dbf4 kinase activity affects specific phosphorylation sites on MCM2 in cancer cells. Journal of cellular biochemistry. 104:1075–1086.

Chawanthayatham, S., C.C. Valentine, 3rd, B.I. Fedeles, E.J. Fox, L.A. Loeb, S.S. Levine, S.L. Slocum, G.N. Wogan, R.G. Croy, and J.M. Essigmann. 2017. Mutational spectra of aflatoxin B1 in vivo establish biomarkers of exposure for human hepatocellular carcinoma. Proceedings of the National Academy of Sciences of the United States of America. 114:E3101–E3109.

Coquerelle, T., J. Dosch, and B. Kaina. 1995. Overexpression of N-methylpurine-DNA glycosylase in Chinese hamster ovary cells renders them more sensitive to the production of chromosomal aberrations by methylating agents--a case of imbalanced DNA repair. Mutation research. 336:9–17.

Cortez, D., G. Glick, and S.J. Elledge. 2004. Minichromosome maintenance proteins are direct targets of the ATM and ATR checkpoint kinases. Proceedings of the National Academy of Sciences of the United States of America. 101:10078–10083.

Crosbie, P.A., A.J. Watson, R. Agius, P.V. Barber, G.P. Margison, and A.C. Povey. 2012. Elevated N3-methylpurine-DNA glycosylase DNA repair activity is associated with lung cancer. Mutation research. 732:43–46.

Cui, F., M.V. Sirotin, and V.B. Zhurkin. 2011. Impact of Alu repeats on the evolution of human p53 binding sites. Biol Direct. 6:2.

Daniel, J.A., M. Pellegrini, J.H. Lee, T.T. Paull, L. Feigenbaum, and A. Nussenzweig. 2008. Multiple autophosphorylation sites are dispensable for murine ATM activation in vivo. The Journal of cell biology. 183:777–783.

Dass, S.B., T.J. Bucci, R.H. Heflich, and D.A. Casciano. 1999. Evaluation of the transgenic p53+/-mouse for detecting genotoxic liver carcinogens in a short-term bioassay. Cancer letters. 143:81–85.

Dass, S.B., G.J. Hammons, T.J. Bucci, R.H. Heflich, and D.A. Casciano. 1998. Susceptibility of C57BL/6 mice to tumorigenicity induced by dimethylnitrosamine and 2-amino-1-methyl-6-phenylimidazo [4,5-b]pyridine in the neonatal bioassay. Cancer letters. 124:105–110.

Dittmann, A., N.J. Kennedy, N.L. Soltero, N. Morshed, M.D. Mana, O.H. Yilmaz, R.J. Davis, and F.M. White. 2019. High-fat diet in a mouse insulin-resistant model induces widespread rewiring of the phosphotyrosine signaling network. Molecular systems biology. 15:e8849.

Doublie, S., and T. Ellenberger. 1998. The mechanism of action of T7 DNA polymerase. Curr Opin Struct Biol. 8:704–712.

Ebrahimkhani, M.R., A. Daneshmand, A. Mazumder, M. Allocca, J.A. Calvo, N. Abolhassani, I. Jhun, S. Muthupalani, C. Ayata, and L.D. Samson. 2014. Aag-initiated base excision repair promotes ischemia reperfusion injury in liver, brain, and kidney. Proceedings of the National Academy of Sciences of the United States of America. 111:E4878–4886.

Engelward, B.P., J.M. Allan, A.J. Dreslin, J.D. Kelly, M.M. Wu, B. Gold, and L.D. Samson. 1998. A chemical and genetic approach together define the biological consequences of 3-methyladenine lesions in the mammalian genome. J Biol Chem. 273:5412–5418.

Engelward, B.P., A. Dreslin, J. Christensen, D. Huszar, C. Kurahara, and L. Samson. 1996. Repair-deficient 3-methyladenine DNA glycosylase homozygous mutant mouse cells have increased sensitivity to alkylation-induced chromosome damage and cell killing. EMBO J. 15:945–952.

Engelward, B.P., G. Weeda, M.D. Wyatt, J.L. Broekhof, J. de Wit, I. Donker, J.M. Allan, B. Gold, J.H. Hoeijmakers, and L.D. Samson. 1997. Base excision repair deficient mice lacking the Aag alkyladenine DNA glycosylase. Proceedings of the National Academy of Sciences of the United States of America. 94:13087–13092.

Ensminger, M., L. Iloff, C. Ebel, T. Nikolova, B. Kaina, and M. Lbrich. 2014. DNA breaks and chromosomal aberrations arise when replication meets base excision repair. The Journal of cell biology. 206:29–43.

Ewald, B., D. Sampath, and W. Plunkett. 2007. H2AX phosphorylation marks gemcitabine-induced stalled replication forks and their collapse upon S-phase checkpoint abrogation. Mol Cancer Ther. 6:1239–1248.

Fahrer, J., J. Frisch, G. Nagel, A. Kraus, B. Dorsam, A.D. Thomas, S. Reissig, A. Waisman, and B. Kaina. 2015. DNA repair by MGMT, but not AAG, causes a threshold in alkylation-induced colorectal carcinogenesis. Carcinogenesis. 36:1235–1244.

Foster, S.S., S. De, L.K. Johnson, J.H. Petrini, and T.H. Stracker. 2012. Cell cycle- and DNA repair pathway-specific effects of apoptosis on tumor suppression. Proceedings of the National Academy of Sciences of the United States of America. 109:9953–9958.

Gatei, M., B. Jakob, P. Chen, A.W. Kijas, O.J. Becherel, N. Gueven, G. Birrell, J.H. Lee, T.T. Paull, Y. Lerenthal, S. Fazry, G. Taucher-Scholz, R. Kalb, D. Schindler, R. Waltes, T. Dork, and M.F. Lavin. 2011. ATM protein-dependent phosphorylation of Rad50 protein regulates DNA repair and cell cycle control. The Journal of biological chemistry. 286:31542–31556.

George, J., M. Tsuchishima, and M. Tsutsumi. 2019. Molecular mechanisms in the pathogenesis of N-nitrosodimethylamine induced hepatic fibrosis. Cell death & disease. 10:18.

Gupta, P.K., A. Sahota, S.A. Boyadjiev, S. Bye, C. Shao, J.P. O’Neill, T.C. Hunter, R.J. Albertini, P.J. Stambrook, and J.A. Tischfield. 1997. High frequency in vivo loss of heterozygosity is primarily a consequence of mitotic recombination. Cancer research. 57:1188–1193.

Haggerty, H.G., and M.P. Holsapple. 1990. Role of metabolism in dimethylnitrosamine-induced immunosuppression: a review. Toxicology. 63:1–23.

Haigis, K.M., and W.F. Dove. 2003. A Robertsonian translocation suppresses a somatic recombination pathway to loss of heterozygosity. Nature genetics. 33:33–39.

Haince, J.F., D. McDonald, A. Rodrigue, U. Dery, J.Y. Masson, M.J. Hendzel, and G.G. Poirier 2008. PARP1-dependent kinetics of recruitment of MRE11 and NBS1 proteins to multiple DNA damage sites. J Biol Chem. 283:1197–1208.

Hall, J., H. Bresil, F. Donato, C.P. Wild, N.A. Loktionova, O.I. Kazanova, I.P. Komyakov, V.G. Lemekhov, A.J. Likhachev, and R. Montesano. 1993. Alkylation and oxidative-DNA damage repair activity in blood leukocytes of smokers and non-smokers. International journal of cancer. Journal international du cancer. 54:728–733.

Hanada, K., M. Budzowska, S.L. Davies, E. van Drunen, H. Onizawa, H.B. Beverloo, A. Maas, J. Essers, I.D. Hickson, and R. Kanaar. 2007. The structure-specific endonuclease Mus81 contributes to replication restart by generating double-strand DNA breaks. Nat Struct Mol Biol. 14:1096–1104.

Hanada, K., M. Budzowska, M. Modesti, A. Maas, C. Wyman, J. Essers, and R. Kanaar. 2006. The structure-specific endonuclease Mus81-Eme1 promotes conversion of interstrand DNA crosslinks into double-strands breaks. The EMBO journal. 25:4921–4932.

Haracska, L., I. Unk, R.E. Johnson, E. Johansson, P.M. Burgers, S. Prakash, and L. Prakash. 2001. Roles of yeast DNA polymerases delta and zeta and of Rev1 in the bypass of abasic sites. Genes & development. 15:945–954.

Hassan, M.M., G. Botrus, R. Abdel-Wahab, R.A. Wolff, D. Li, D. Tweardy, A.T. Phan, E. Hawk, M. Javle, J.S. Lee, H.A. Torres, A. Rashid, R. Lenzi, H.M. Hassabo, Y. Abaza, A.S. Shalaby, S. Lacin, J. Morris, Y.Z. Patt, C.I. Amos, S.A. Khaderi, J.A. Goss, P.K. Jalal, and A.O. Kaseb. 2017. Estrogen Replacement Reduces Risk and Increases Survival Times of Women With Hepatocellular Carcinoma. Clin Gastroenterol Hepatol. 15:1791–1799.

He, M., R.G. Croy, J.M. Essigmann, and T.M. Swager. 2019. Chemiresistive Carbon Nanotube Sensors for N-Nitrosodialkylamines. ACS Sens.

Heath, D.F.M., A. R.. 1961. Preparation of 14C-labelled dialkylnitrosamines and an improved preparation of N-methyl-N-t-butylamine. J Chem Soc:4226–4229.

Hendricks, C.A., M. Razlog, T. Matsuguchi, A. Goyal, A.L. Brock, and B.P. Engelward. 2002. The S. cerevisiae Mag1 3-methyladenine DNA glycosylase modulates susceptibility to homologous recombination. DNA repair. 1:645–659.

Huang, X., M. Okafuji, F. Traganos, E. Luther, E. Holden, and Z. Darzynkiewicz. 2004. Assessment of histone H2AX phosphorylation induced by DNA topoisomerase I and II inhibitors topotecan and mitoxantrone and by the DNA cross-linking agent cisplatin. Cytometry. Part A : the journal of the International Society for Analytical Cytology. 58:99–110.

Ibeanu, G., B. Hartenstein, W.C. Dunn, L.Y. Chang, E. Hofmann, T. Coquerelle, S. Mitra, and B. Kaina. 1992. Overexpression of human DNA repair protein N-methylpurine-DNA glycosylase results in the increased removal of N-methylpurines in DNA without a concomitant increase in resistance to alkylating agents in Chinese hamster ovary cells. Carcinogenesis. 13:1989–1995.

Iyer, S.S., W.P. Pulskens, J.J. Sadler, L.M. Butter, G.J. Teske, T.K. Ulland, S.C. Eisenbarth, S. Florquin, R.A. Flavell, J.C. Leemans, and F.S. Sutterwala. 2009. Necrotic cells trigger a sterile inflammatory response through the Nlrp3 inflammasome. Proceedings of the National Academy of Sciences of the United States of America. 106:20388–20393.

Johnson, R.E., S.L. Yu, S. Prakash, and L. Prakash. 2007. A role for yeast and human translesion synthesis DNA polymerases in promoting replication through 3-methyl adenine. Molecular and cellular biology. 27:7198–7205.

Kaina, B., G. Fritz, and T. Coquerelle. 1993. Contribution of O6-alkylguanine and N-alkylpurines to the formation of sister chromatid exchanges, chromosomal aberrations, and gene mutations: new insights gained from studies of genetically engineered mammalian cell lines. Environmental and molecular mutagenesis. 22:283–292.

Karabatsos, G.J.T., R. A. 1964. Structural studies by Nuclear Magnetic Resonance. IX. Configuration of N-Nitrosamines. Journal of the American Chemical Society. 86:4373–4378.

Kimoto, W. 1980. Role of strong ion exchange resins in nitrosamine formation in water. Water Res.

Kiraly, O., G. Gong, M.D. Roytman, Y. Yamada, L.D. Samson, and B.P. Engelward. 2014. DNA glycosylase activity and cell proliferation are key factors in modulating homologous recombination in vivo. Carcinogenesis. 35:2495–2502.

Kisby, G.E., A. Olivas, T. Park, M. Churchwell, D. Doerge, L.D. Samson, S.L. Gerson, and M.S. Turker. 2009. DNA repair modulates the vulnerability of the developing brain to alkylating agents. DNA repair. 8:400–412.

Klapacz, J., G.M. Lingaraju, H.H. Guo, D. Shah, A. Moar-Shoshani, L.A. Loeb, and L.D. Samson. 2010. Frameshift mutagenesis and microsatellite instability induced by human alkyladenine DNA glycosylase. Molecular cell. 37:843–853.

Kolomietz, E., M.S. Meyn, A. Pandita, and J.A. Squire. 2002. The role of Alu repeat clusters as mediators of recurrent chromosomal aberrations in tumors. Genes Chromosomes Cancer. 35:97–112.

Krokan, H.E., and M. Bjoras. 2013. Base excision repair. Cold Spring Harb Perspect Biol. 5:a012583.

Lambert, I.B., T.M. Singer, S.E. Boucher, and G.R. Douglas. 2005. Detailed review of transgenic rodent mutation assays. Mutation research. 590:1–280.

Lambert, S., K. Mizuno, J. Blaisonneau, S. Martineau, R. Chanet, K. Freon, J.M. Murray, A.M. Carr, and G. Baldacci. 2010. Homologous recombination restarts blocked replication forks at the expense of genome rearrangements by template exchange. Molecular cell. 39:346–359.

Larson, K., J. Sahm, R. Shenkar, and B. Strauss. 1985. Methylation-induced blocks to in vitro DNA replication. Mutation research. 150:77–84.

Lee, V.M., L.K. Keefer, and M.C. Archer. 1996. An evaluation of the roles of metabolic denitrosation and alpha-hydroxylation in the hepatotoxicity of N-Nitrosodimethylamine. Chem Res Toxicol. 9:1319–1324.

Leitner-Dagan, Y., Z. Sevilya, M. Pinchev, R. Kramer, D. Elinger, L.C. Roisman, H.S. Rennert, E. Schechtman, L. Freedman, G. Rennert, Z. Livneh, and T. Paz-Elizur. 2012. N-methylpurine DNA glycosylase and OGG1 DNA repair activities: opposite associations with lung cancer risk. J Natl Cancer Inst. 104:1765–1769.

Li, M., and X. Yu. 2013. Function of BRCA1 in the DNA damage response is mediated by ADP-ribosylation. Cancer cell. 23:693–704.

Li, Q., Q. Hao, W. Cao, J. Li, K. Wu, Y. Elshimali, D. Zhu, Q.H. Chen, G. Chen, J.R. Pollack, J. Vadgama, and Y. Wu. 2019. PP2Cdelta inhibits p300-mediated p53 acetylation via ATM/BRCA1 pathway to impede DNA damage response in breast cancer. Sci Adv. 5:eaaw8417.

Likhite, V.S., E.I. Cass, S.D. Anderson, J.R. Yates, and A.M. Nardulli. 2004. Interaction of estrogen receptor alpha with 3-methyladenine DNA glycosylase modulates transcription and DNA repair. The Journal of biological chemistry. 279:16875–16882.

Loktionov, A., M. Hollstein, N. Martel, D. Galendo, J.R. Cabral, L. Tomatis, and H. Yamasaki. 1990. Tissue-specific activating mutations of Ha- and Ki-ras oncogenes in skin, lung, and liver tumors induced in mice following transplacental exposure to DMBA. Mol Carcinog. 3:134–140.

Lowe, S.W., and A.W. Lin. 2000. Apoptosis in cancer. Carcinogenesis. 21:485–495.

Margulies, C.M., I.A. Chaim, A. Mazumder, J. Criscione, and L.D. Samson. 2017. Alkylation induced cerebellar degeneration dependent on Aag and Parp1 does not occur via previously established cell death mechanisms. PloS one. 12:e0184619.

Marians, K.J. 2018. Lesion Bypass and the Reactivation of Stalled Replication Forks. Annu Rev Biochem. 87:217–238.

Matsuoka, S., B.A. Ballif, A. Smogorzewska, E.R. McDonald, 3rd, K.E. Hurov, J. Luo, C.E. Bakalarski, Z. Zhao, N. Solimini, Y. Lerenthal, Y. Shiloh, S.P. Gygi, and S.J. Elledge. 2007. ATM and ATR substrate analysis reveals extensive protein networks responsive to DNA damage. Science. 316:1160–1166.

Mazouzi, A., A. Stukalov, A.C. Muller, D. Chen, M. Wiedner, J. Prochazkova, S.C. Chiang, M. Schuster, F.P. Breitwieser, A. Pichlmair, S.F. El-Khamisy, C. Bock, R. Kralovics, J. Colinge, K.L. Bennett, and J.I. Loizou. 2016. A Comprehensive Analysis of the Dynamic Response to Aphidicolin-Mediated Replication Stress Uncovers Targets for ATM and ATMIN. Cell Rep. 15:893–908.

Meira, L.B., J.M. Bugni, S.L. Green, C.W. Lee, B. Pang, D. Borenshtein, B.H. Rickman, A.B. Rogers, C.A. Moroski-Erkul, J.L. McFaline, D.B. Schauer, P.C. Dedon, J.G. Fox, and L.D. Samson. 2008. DNA damage induced by chronic inflammation contributes to colon carcinogenesis in mice. The Journal of clinical investigation. 118:2516–2525.

Meira, L.B., C.A. Moroski-Erkul, S.L. Green, J.A. Calvo, R.T. Bronson, D. Shah, and L.D. Samson. 2009. Aag-initiated base excision repair drives alkylation-induced retinal degeneration in mice. Proceedings of the National Academy of Sciences of the United States of America. 106:888–893.

Mientjes, E.J., A. Luiten-Schuite, E. van der Wolf, Y. Borsboom, A. Bergmans, F. Berends, P.H. Lohman, R.A. Baan, and J.H. van Delft. 1998. DNA adducts, mutant frequencies, and mutation spectra in various organs of lambda lacZ mice exposed to ethylating agents. Environmental and molecular mutagenesis. 31:18–31.

Mitch, W.A., and D.L. Sedlak. 2004. Characterization and fate of N-nitrosodimethylamine precursors in municipal wastewater treatment plants. Environ Sci Technol. 38:1445–1454.

Moeglin, E., D. Desplancq, S. Conic, M. Oulad-Abdelghani, A. Stoessel, M. Chiper, M. Vigneron, P. Didier, L. Tora, and E. Weiss. 2019. Uniform Widespread Nuclear Phosphorylation of Histone H2AX Is an Indicator of Lethal DNA Replication Stress. Cancers (Basel). 11.

Monti, P., I. Traverso, L. Casolari, P. Menichini, A. Inga, L. Ottaggio, D. Russo, P. Iyer, B. Gold, and G. Fronza. 2010. Mutagenicity of N3-methyladenine: a multi-translesion polymerase affair. Mutation research. 683:50–56.

Nagel, Z.D., C.M. Margulies, I.A. Chaim, S.K. McRee, P. Mazzucato, A. Ahmad, R.P. Abo, V.L. Butty, A.L. Forget, and L.D. Samson. 2014. Multiplexed DNA repair assays for multiple lesions and multiple doses via transcription inhibition and transcriptional mutagenesis. Proceedings of the National Academy of Sciences of the United States of America. 111:E1823–1832.

Naugler, W.E., T. Sakurai, S. Kim, S. Maeda, K. Kim, A.M. Elsharkawy, and M. Karin. 2007. Gender disparity in liver cancer due to sex differences in MyD88-dependent IL-6 production. Science. 317:121–124.

Nishikawa, A., F. Furukawa, K. Kasahara, I.S. Lee, T. Suzuki, M. Hayashi, T. Sofuni, and M. Takahashi. 1997. Comparative study on organ-specificity of tumorigenicity, mutagenicity and cell proliferative activity induced by dimethylnitrosamine in Big Blue mice. Cancer letters. 117:143–147.

Nohmi, T., M. Katoh, H. Suzuki, M. Matsui, M. Yamada, M. Watanabe, M. Suzuki, N. Horiya, O. Ueda, T. Shibuya, H. Ikeda, and T. Sofuni. 1996. A new transgenic mouse mutagenesis test system using Spi- and 6-thioguanine selections. Environmental and molecular mutagenesis. 28:465–470.

Norbury, C.J., and B. Zhivotovsky. 2004. DNA damage-induced apoptosis. Oncogene. 23:2797–2808.

Ogiwara, H., T. Kohno, H. Nakanishi, K. Nagayama, M. Sato, and J. Yokota. 2008. Unbalanced translocation, a major chromosome alteration causing loss of heterozygosity in human lung cancer. Oncogene. 27:4788–4797.

Pages, V., R.E. Johnson, L. Prakash, and S. Prakash. 2008. Mutational specificity and genetic control of replicative bypass of an abasic site in yeast. Proceedings of the National Academy of Sciences of the United States of America. 105:1170–1175.

Pal, J., R. Bertheau, L. Buon, A. Qazi, R.B. Batchu, S. Bandyopadhyay, R. Ali-Fehmi, D.G. Beer, D.W. Weaver, R.J. Shmookler Reis, R.K. Goyal, Q. Huang, N.C. Munshi, and M.A. Shammas. 2011. Genomic evolution in Barrett’s adenocarcinoma cells: critical roles of elevated hsRAD51, homologous recombination and Alu sequences in the genome. Oncogene. 30:3585–3598.

Parr, M.K., and J.F. Joseph. 2019. NDMA impurity in valsartan and other pharmaceutical products: Analytical methods for the determination of N-nitrosamines. J Pharm Biomed Anal. 164:536–549.

Parrish, M.C., I.A. Chaim, Z.D. Nagel, S.R. Tannenbaum, L.D. Samson, and B.P. Engelward. 2018. Nitric oxide induced S-nitrosation causes base excision repair imbalance. DNA repair. 68:25–33.

Pegg, A.E., and G. Hui. 1978. Formation and subsequent removal of O6-methylguanine from deoxyribonucleic acid in rat liver and kidney after small doses of dimethylnitrosamine. The Biochemical journal. 173:739–748.

Pellegrini, M., A. Celeste, S. Difilippantonio, R. Guo, W. Wang, L. Feigenbaum, and A. Nussenzweig. 2006. Autophosphorylation at serine 1987 is dispensable for murine Atm activation in vivo. Nature. 443:222–225.

Perez-Riverol, Y., A. Csordas, J. Bai, M. Bernal-Llinares, S. Hewapathirana, D.J. Kundu, A. Inuganti, J. Griss, G. Mayer, M. Eisenacher, E. Perez, J. Uszkoreit, J. Pfeuffer, T. Sachsenberg, S. Yilmaz, S. Tiwary, J. Cox, E. Audain, M. Walzer, A.F. Jarnuczak, T. Ternent, A. Brazma, and J.A. Vizcaino. 2019. The PRIDE database and related tools and resources in 2019: improving support for quantification data. Nucleic acids research. 47:D442–D450.

Peto, R., R. Gray, P. Brantom, and P. Grasso. 1991. Dose and time relationships for tumor induction in the liver and esophagus of 4080 inbred rats by chronic ingestion of N-nitrosodiethylamine or N-nitrosodimethylamine. Cancer research. 51:6452–6469.

Piazza, A., W.D. Wright, and W.D. Heyer. 2017. Multi-invasions Are Recombination Byproducts that Induce Chromosomal Rearrangements. Cell. 170:760–773 e715.

Plosky, B.S., E.G. Frank, D.A. Berry, G.P. Vennall, J.P. McDonald, and R. Woodgate. 2008. Eukaryotic Y-family polymerases bypass a 3-methyl-2’-deoxyadenosine analog in vitro and methyl methanesulfonate-induced DNA damage in vivo. Nucleic acids research. 36:2152–2162.

Pottegard, A., K.B. Kristensen, M.T. Ernst, N.B. Johansen, P. Quartarolo, and J. Hallas. 2018. Use of N-nitrosodimethylamine (NDMA) contaminated valsartan products and risk of cancer: Danish nationwide cohort study. Bmj. 362:k3851.

Rao, K.V., and S.D. Vesselinovitch. 1973. Age- and sex-associated diethylnitrosamine dealkylation activity of the mouse liver and hepatocarcinogenesis. Cancer research. 33:1625–1627.

Richardson, S.D. 2003. Disinfection by-products and other emerging contaminants in drinking water. Trends in Analytical Chemistry.

Robertson, A.B., A. Klungland, T. Rognes, and I. Leiros. 2009. DNA repair in mammalian cells: Base excision repair: the long and short of it. Cell Mol Life Sci. 66:981–993.

Roos, W.P., and B. Kaina. 2013. DNA damage-induced cell death: from specific DNA lesions to the DNA damage response and apoptosis. Cancer letters. 332:237–248.

Sachet, M., Y.Y. Liang, and R. Oehler. 2017. The immune response to secondary necrotic cells. Apoptosis. 22:1189–1204.

Saugar, I., M.V. Vazquez, M. Gallo-Fernandez, M.A. Ortiz-Bazan, M. Segurado, A. Calzada, and J.A. Tercero. 2013. Temporal regulation of the Mus81-Mms4 endonuclease ensures cell survival under conditions of DNA damage. Nucleic acids research. 41:8943–8958.

Scherf-Clavel, O., M. Kinzig, A. Besa, A. Schreiber, C. Bidmon, M. Abdel-Tawab, J. Wohlfart, F. Sorgel, and U. Holzgrabe. 2019. The contamination of valsartan and other sartans, Part 2: Untargeted screening reveals contamination with amides additionally to known nitrosamine impurities. J Pharm Biomed Anal. 172:278–284.

Schmezer, P., C. Eckert, and U.M. Liegibel. 1994. Tissue-specific induction of mutations by streptozotocin in vivo. Mutation research. 307:495–499.

Scully, R., and A. Xie. 2013. Double strand break repair functions of histone H2AX. Mutation research. 750:5–14.

Sedlak, D.L., R.A. Deeb, E.L. Hawley, W.A. Mitch, T.D. Durbin, S. Mowbray, and S. Carr. 2005. Sources and fate of nitrosodimethylamine and its precursors in municipal wastewater treatment plants. Water Environ Res. 77:32–39.

Shao, C., L. Deng, O. Henegariu, L. Liang, N. Raikwar, A. Sahota, P.J. Stambrook, and J.A. Tischfield. 1999. Mitotic recombination produces the majority of recessive fibroblast variants in heterozygous mice. Proceedings of the National Academy of Sciences of the United States of America. 96:9230–9235.

Snider, T.A., A. Richardson, J.A. Stoner, and S.S. Deepa. 2018. The Geropathology Grading Platform demonstrates that mice null for Cu/Zn-superoxide dismutase show accelerated biological aging. Geroscience. 40:97–103.

Sobol, R.W., M. Kartalou, K.H. Almeida, D.F. Joyce, B.P. Engelward, J.K. Horton, R. Prasad, L.D. Samson, and S.H. Wilson. 2003. Base excision repair intermediates induce p53-independent cytotoxic and genotoxic responses. J Biol Chem. 278:39951–39959.

Sohn, O.S., H. Ishizaki, C.S. Yang, and E.S. Fiala. 1991. Metabolism of azoxymethane, methylazoxymethanol and N-nitrosodimethylamine by cytochrome P450IIE1. Carcinogenesis. 12:127–131.

Sorgel, F., M. Kinzig, M. Abdel-Tawab, C. Bidmon, A. Schreiber, S. Ermel, J. Wohlfart, A. Besa, O. Scherf-Clavel, and U. Holzgrabe. 2019. The contamination of valsartan and other sartans, part 1: New findings. J Pharm Biomed Anal. 172:395–405.

Soriano, P. 1999. Generalized lacZ expression with the ROSA26 Cre reporter strain. Nature genetics. 21:70–71.

Souliotis, V.L., J.H. van Delft, M.J. Steenwinkel, R.A. Baan, and S.A. Kyrtopoulos. 1998. DNA adducts, mutant frequencies and mutation spectra in lambda lacZ transgenic mice treated with N-nitrosodimethylamine. Carcinogenesis. 19:731–739.

Strauss, B., D. Scudiero, and E. Henderson. 1975. The nature of the alkylation lesion in mammalian cells. Basic Life Sci. 5A:13–24.

Strout, M.P., G. Marcucci, C.D. Bloomfield, and M.A. Caligiuri. 1998. The partial tandem duplication of ALL1 (MLL) is consistently generated by Alu-mediated homologous recombination in acute myeloid leukemia. Proceedings of the National Academy of Sciences of the United States of America. 95:2390–2395.

Stucki, M., J.A. Clapperton, D. Mohammad, M.B. Yaffe, S.J. Smerdon, and S.P. Jackson. 2005. MDC1 directly binds phosphorylated histone H2AX to regulate cellular responses to DNA double-strand breaks. Cell. 123:1213–1226.

Sukup-Jackson, M.R., O. Kiraly, J.E. Kay, L. Na, E.A. Rowland, K.E. Winther, D.N. Chow, T. Kimoto, T. Matsuguchi, V.S. Jonnalagadda, V.I. Maklakova, V.R. Singh, D.N. Wadduwage, J. Rajapakse, P.T. So, L.S. Collier, and B.P. Engelward. 2014. Rosa26-GFP direct repeat (RaDR-GFP) mice reveal tissue- and age-dependence of homologous recombination in mammals in vivo. PLoS genetics. 10:e1004299.

Swenberg, J.A., D.G. Hoel, and P.N. Magee. 1991. Mechanistic and statistical insight into the large carcinogenesis bioassays on N-nitrosodiethylamine and N-nitrosodimethylamine. Cancer research. 51:6409–6414.

Taus, T., T. Kocher, P. Pichler, C. Paschke, A. Schmidt, C. Henrich, and K. Mechtler. 2011. Universal and confident phosphorylation site localization using phosphoRS. J Proteome Res. 10:5354–5362.

Thoolen, B., R.R. Maronpot, T. Harada, A. Nyska, C. Rousseaux, T. Nolte, D.E. Malarkey, W. Kaufmann, K. Kuttler, U. Deschl, D. Nakae, R. Gregson, M.P. Vinlove, A.E. Brix, B. Singh, F. Belpoggi, and J.M. Ward. 2010. Proliferative and nonproliferative lesions of the rat and mouse hepatobiliary system. Toxicologic pathology. 38:5S–81S.

Tolba, R., T. Kraus, C. Liedtke, M. Schwarz, and R. Weiskirchen. 2015. Diethylnitrosamine (DEN)-induced carcinogenic liver injury in mice. Lab Anim. 49:59–69.

Tsutsumi, T., H. Akiyama, Y. Demizu, N. Uchiyama, S. Masada, G. Tsuji, R. Arai, Y. Abe, T. Hakamatsuka, K.I. Izutsu, Y. Goda, and H. Okuda. 2019. Analysis of an Impurity, N-Nitrosodimethylamine, in Valsartan Drug Substances and Associated Products Using GC-MS. Biol Pharm Bull. 42:547–551.

Vesselinovitch, S.D. 1969. The sex-dependent difference in the development of liver tumors in mice administered dimethylnitrosamine. Cancer research. 29:1024–1027.

Wadduwage, D.N., J. Kay, V.R. Singh, O. Kiraly, M.R. Sukup-Jackson, J. Rajapakse, B.P. Engelward, and P.T.C. So. 2018. Automated fluorescence intensity and gradient analysis enables detection of rare fluorescent mutant cells deep within the tissue of RaDR mice. Sci Rep. 8:12108.

Wallace, S.S., D.L. Murphy, and J.B. Sweasy. 2012. Base excision repair and cancer. Cancer letters. 327:73–89.

Wang, B., S. Matsuoka, B.A. Ballif, D. Zhang, A. Smogorzewska, S.P. Gygi, and S.J. Elledge. 2007. Abraxas and RAP80 form a BRCA1 protein complex required for the DNA damage response. Science. 316:1194–1198.

Wang, X., T. Suzuki, T. Itoh, M. Honma, A. Nishikawa, F. Furukawa, M. Takahashi, M. Hayashi, T. Kato, and T. Sofuni. 1998. Specific mutational spectrum of dimethylnitrosamine in the lacI transgene of Big Blue C57BL/6 mice. Mutagenesis. 13:625–630.

Ward, I.M., and J. Chen. 2001. Histone H2AX is phosphorylated in an ATR-dependent manner in response to replicational stress. The Journal of biological chemistry. 276:47759–47762.

Weghorst, C.M., M.A. Pereira, and J.E. Klaunig. 1989. Strain differences in hepatic tumor promotion by phenobarbital in diethylnitrosamine- and dimethylnitrosamine-initiated infant male mice. Carcinogenesis. 10:1409–1412.

Westman, J., S. Grinstein, and P.E. Marques. 2019. Phagocytosis of Necrotic Debris at Sites of Injury and Inflammation. Front Immunol. 10:3030.

Willis, N., and N. Rhind. 2009. Mus81, Rhp51(Rad51), and Rqh1 form an epistatic pathway required for the S-phase DNA damage checkpoint. Molecular biology of the cell. 20:819–833.

Wirtz, S., G. Nagel, L. Eshkind, M.F. Neurath, L.D. Samson, and B. Kaina. 2010. Both base excision repair and O6-methylguanine-DNA methyltransferase protect against methylation-induced colon carcinogenesis. Carcinogenesis. 31:2111–2117.

Xu, B., S. Kim, and M.B. Kastan. 2001. Involvement of Brca1 in S-phase and G(2)-phase checkpoints after ionizing irradiation. Molecular and cellular biology. 21:3445–3450.

Xu, Y., S. Huang, Z.G. Liu, and J. Han. 2006. Poly(ADP-ribose) polymerase-1 signaling to mitochondria in necrotic cell death requires RIP1/TRAF2-mediated JNK1 activation. The Journal of biological chemistry. 281:8788–8795.

Yang, Q., W. Lin, Z. Liu, J. Zhu, N. Huang, Z. Cui, Z. Han, Q. Pan, A. Goel, and F. Sun. 2018. RAP80 is an independent prognosis biomarker for the outcome of patients with esophageal squamous cell carcinoma. Cell death & disease. 9:146.

Yeeles, J.T., J. Poli, K.J. Marians, and P. Pasero. 2013. Rescuing stalled or damaged replication forks. Cold Spring Harb Perspect Biol. 5:a012815.

Yoon, J.H., J. Roy Choudhury, J. Park, S. Prakash, and L. Prakash. 2017. Translesion synthesis DNA polymerases promote error-free replication through the minor-groove DNA adduct 3-deaza-3-methyladenine. The Journal of biological chemistry. 292:18682–18688.

Yu, S.W., H. Wang, M.F. Poitras, C. Coombs, W.J. Bowers, H.J. Federoff, G.G. Poirier, T.M. Dawson, and V.L. Dawson. 2002. Mediation of poly(ADP-ribose) polymerase-1-dependent cell death by apoptosis-inducing factor. Science. 297:259–263.

Zhang, W., A. Edwards, W. Fan, P. Deininger, and K. Zhang. 2011. Alu distribution and mutation types of cancer genes. BMC genomics. 12:157.

Zhang, Y., F. Yuan, X. Wu, O. Rechkoblit, J.S. Taylor, N.E. Geacintov, and Z. Wang. 2000. Error-prone lesion bypass by human DNA polymerase eta. Nucleic acids research. 28:4717–4724.

Zhao, L., H. Lin, S. Chen, S. Chen, M. Cui, D. Shi, B. Wang, K. Ma, and Z. Shao. 2018. Hydrogen peroxide induces programmed necrosis in rat nucleus pulposus cells through the RIP1/RIP3-PARP-AIF pathway. J Orthop Res. 36:1269–1282.

