## Supplemental figures for "Excision of mutagenic replication-blocking lesions suppresses cancer but promotes cytotoxicity and lethality in nitrosamine-exposed mice"

### Slide 1
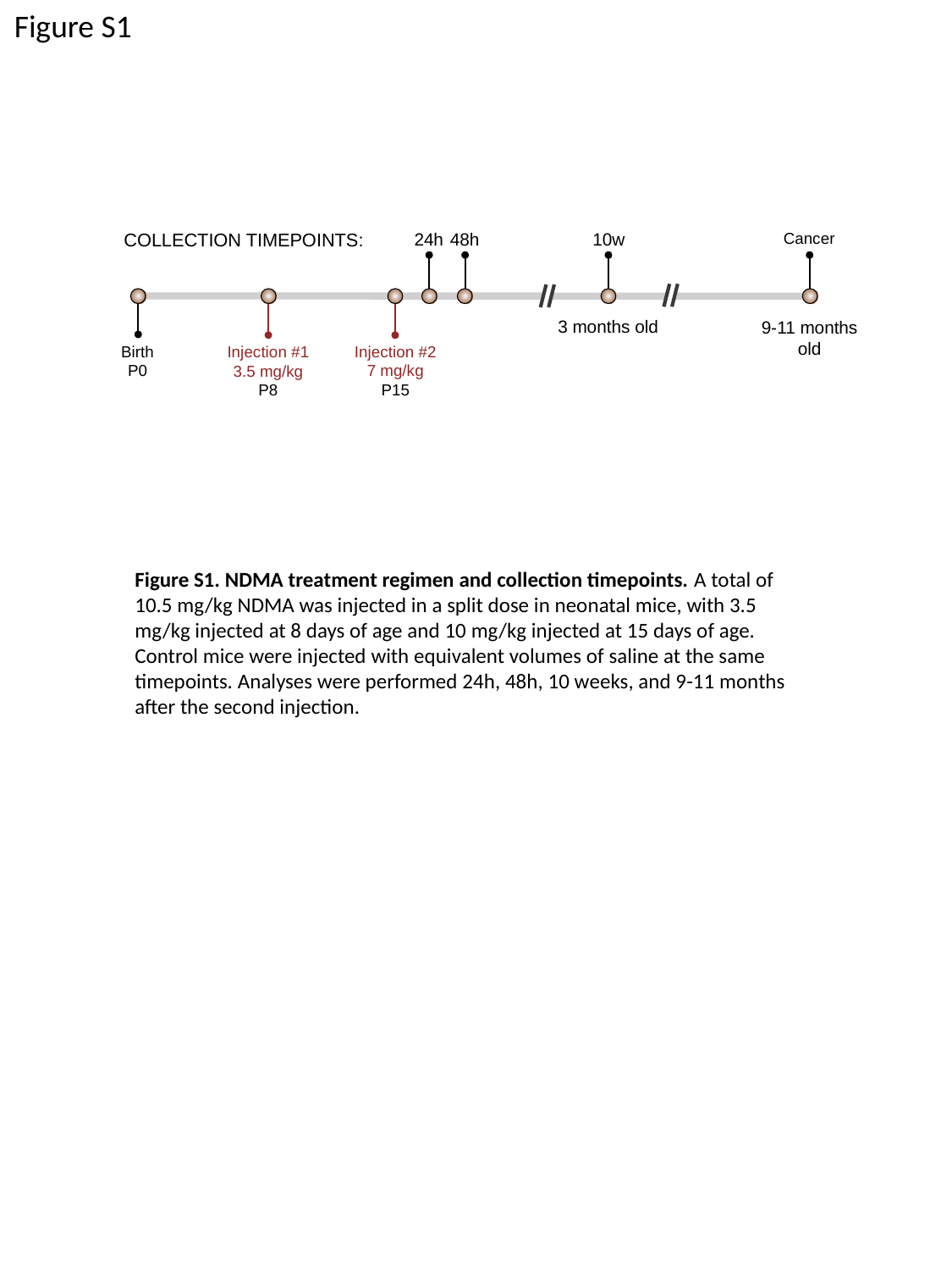

Figure S1
COLLECTION TIMEPOINTS:
24h
48h
10w
Cancer
3 months old
9-11 months old
Birth
P0
Injection #2
7 mg/kg
P15
Injection #1
3.5 mg/kg
P8
Figure S1. NDMA treatment regimen and collection timepoints. A total of 10.5 mg/kg NDMA was injected in a split dose in neonatal mice, with 3.5 mg/kg injected at 8 days of age and 10 mg/kg injected at 15 days of age. Control mice were injected with equivalent volumes of saline at the same timepoints. Analyses were performed 24h, 48h, 10 weeks, and 9-11 months after the second injection.

### Slide 2
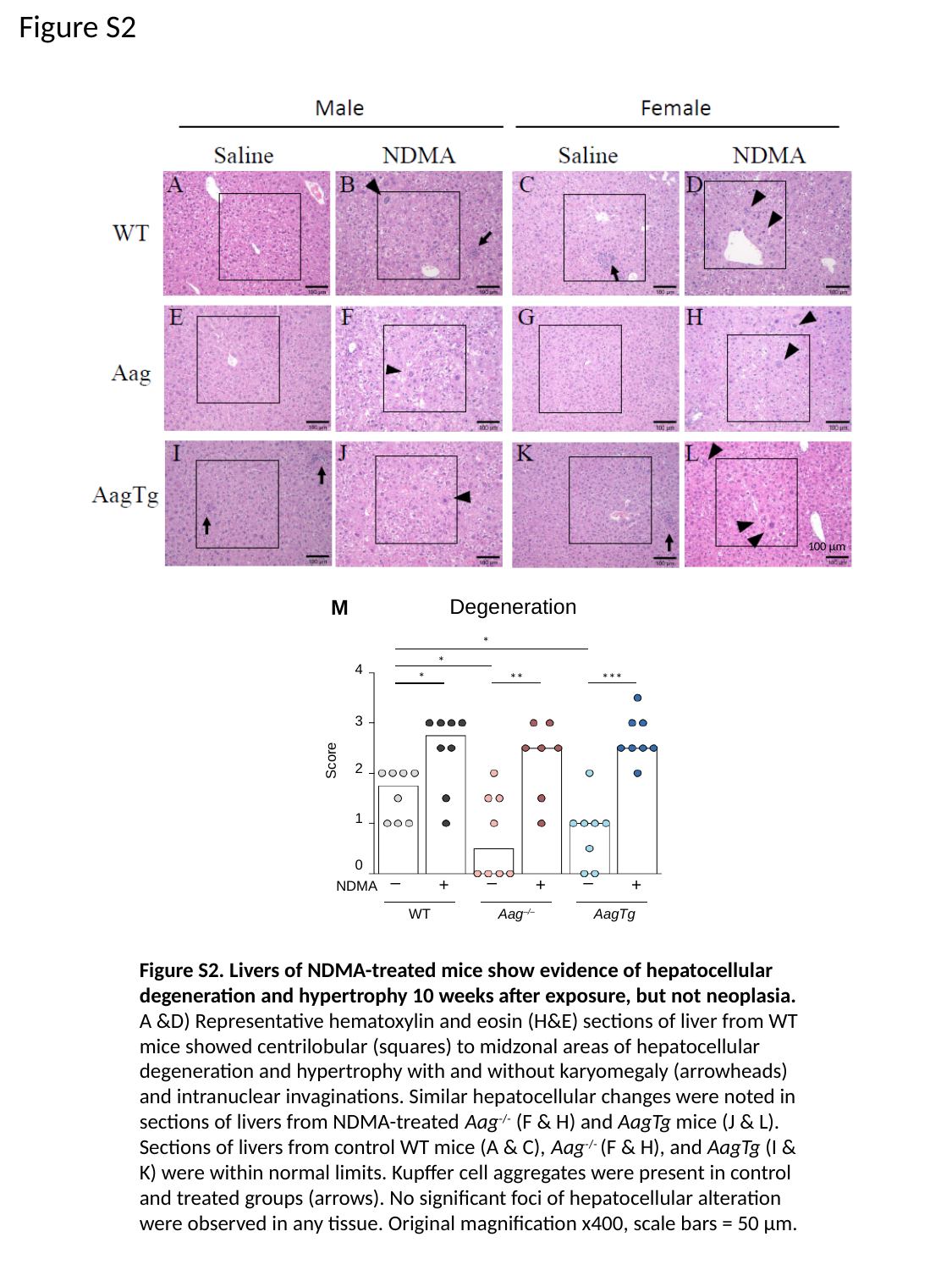

Figure S2
100 μm
Degeneration
M
*
*
4
*
***
**
3
Score
2
1
0
–
–
–
+
+
+
NDMA
AagTg
Aag^–/–^
WT
Figure S2. Livers of NDMA-treated mice show evidence of hepatocellular degeneration and hypertrophy 10 weeks after exposure, but not neoplasia. A &D) Representative hematoxylin and eosin (H&E) sections of liver from WT mice showed centrilobular (squares) to midzonal areas of hepatocellular degeneration and hypertrophy with and without karyomegaly (arrowheads) and intranuclear invaginations. Similar hepatocellular changes were noted in sections of livers from NDMA-treated Aag^-/-^ (F & H) and AagTg mice (J & L). Sections of livers from control WT mice (A & C), Aag^-/-^ (F & H), and AagTg (I & K) were within normal limits. Kupffer cell aggregates were present in control and treated groups (arrows). No significant foci of hepatocellular alteration were observed in any tissue. Original magnification x400, scale bars = 50 μm.

### Slide 3
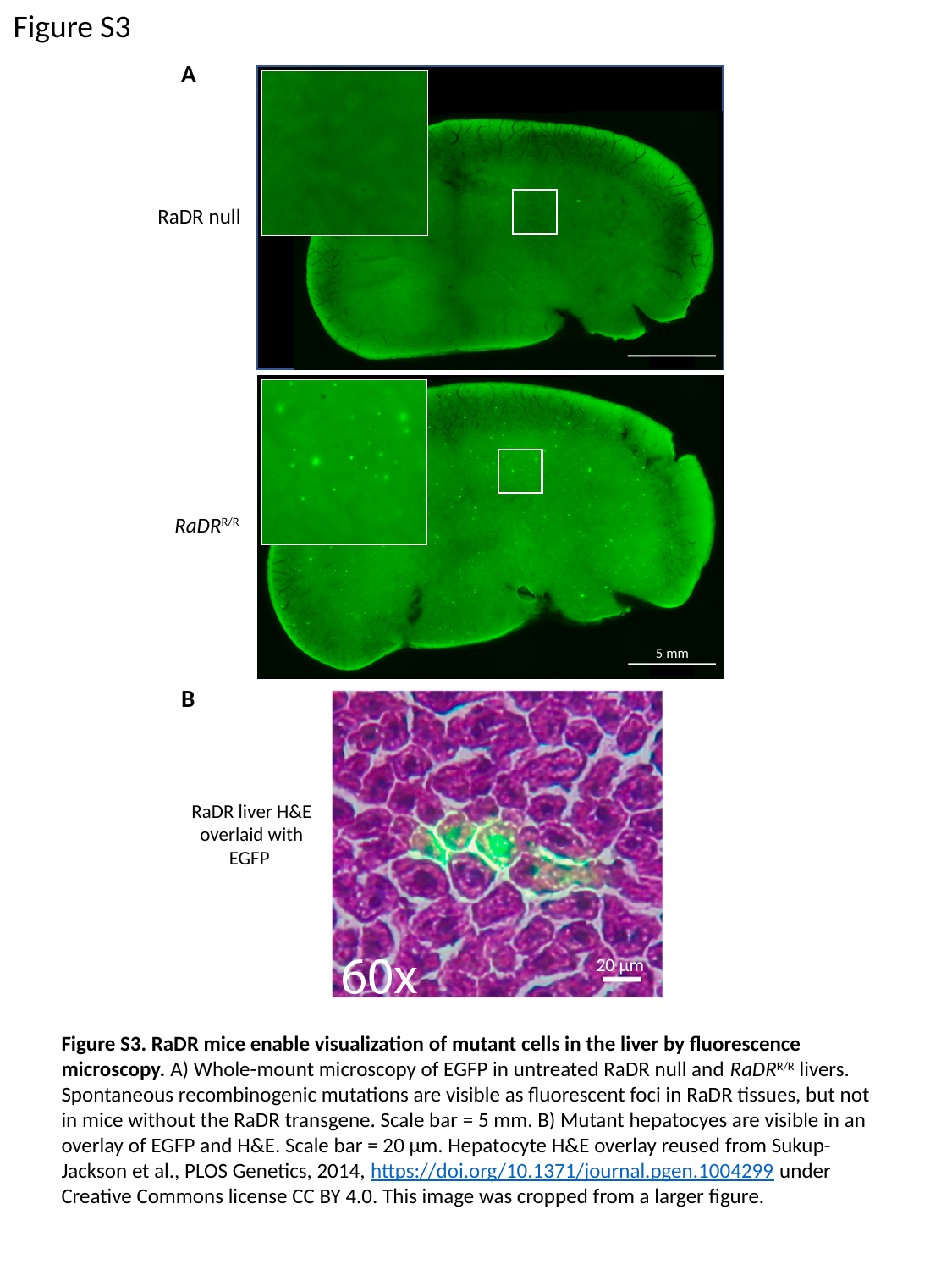

Figure S3
A
RaDR null
RaDR^R/R^
5 mm
B
RaDR liver H&E overlaid with EGFP
20 μm
Figure S3. RaDR mice enable visualization of mutant cells in the liver by fluorescence microscopy. A) Whole-mount microscopy of EGFP in untreated RaDR null and RaDR^R/R^ livers. Spontaneous recombinogenic mutations are visible as fluorescent foci in RaDR tissues, but not in mice without the RaDR transgene. Scale bar = 5 mm. B) Mutant hepatocyes are visible in an overlay of EGFP and H&E. Scale bar = 20 μm. Hepatocyte H&E overlay reused from Sukup-Jackson et al., PLOS Genetics, 2014, https://doi.org/10.1371/journal.pgen.1004299 under Creative Commons license CC BY 4.0. This image was cropped from a larger figure.

### Slide 4
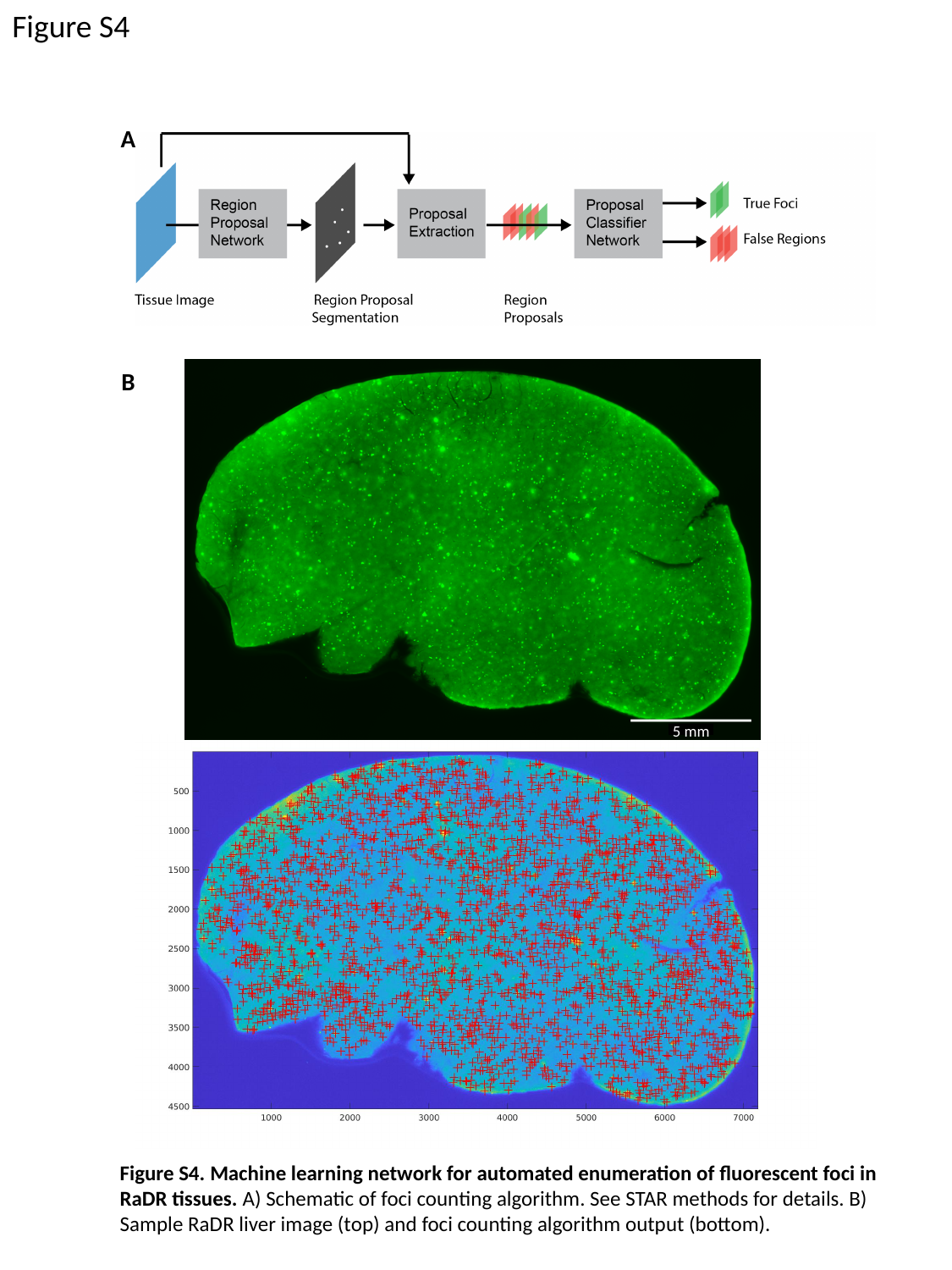

Figure S4
A
B
5 mm
Figure S4. Machine learning network for automated enumeration of fluorescent foci in RaDR tissues. A) Schematic of foci counting algorithm. See STAR methods for details. B) Sample RaDR liver image (top) and foci counting algorithm output (bottom).

### Slide 5
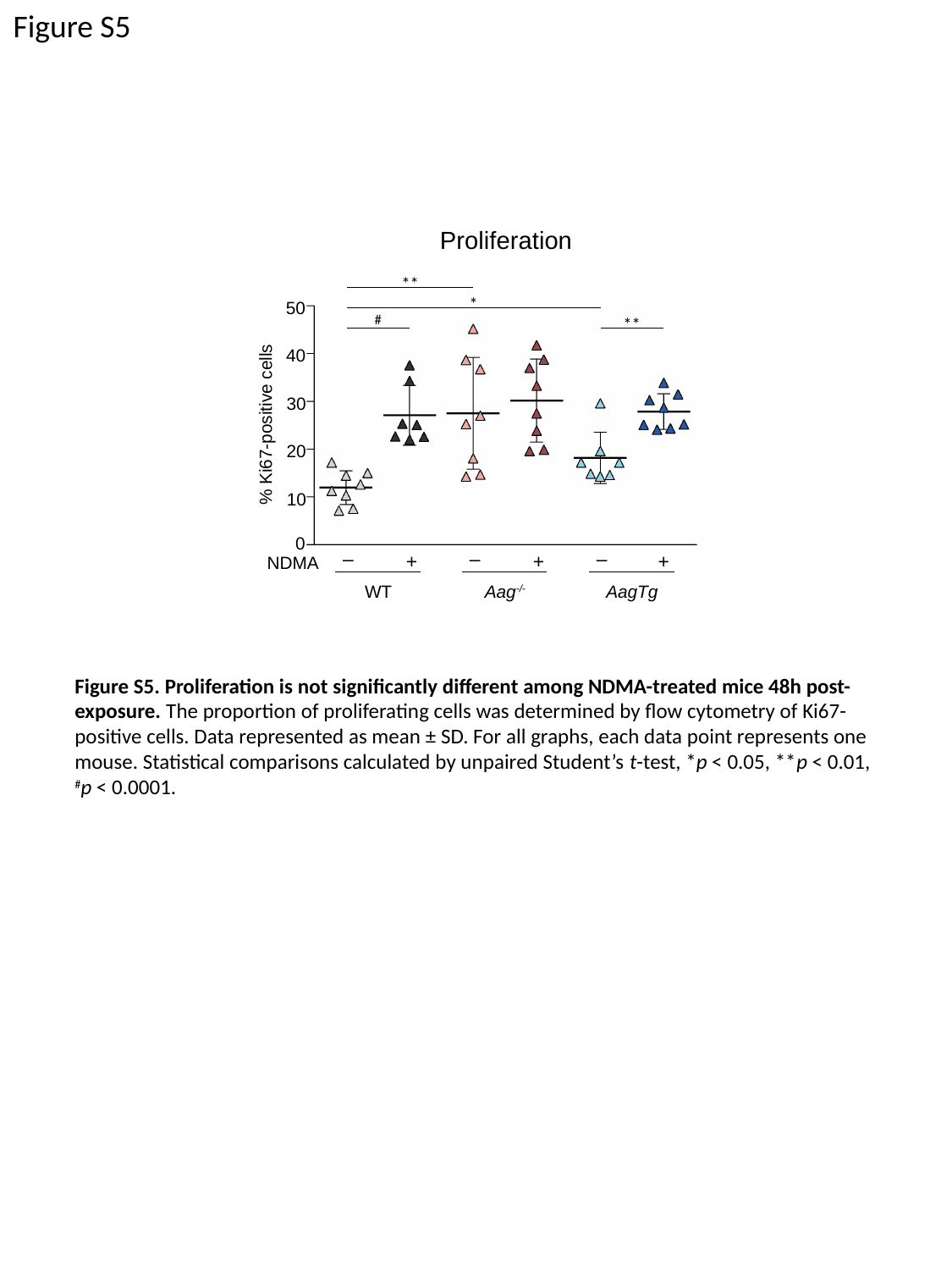

Figure S5
Proliferation
**
*
50
#
**
40
30
% Ki67-positive cells
20
10
0
–
–
–
+
+
+
NDMA
AagTg
Aag^-/-^
WT
Figure S5. Proliferation is not significantly different among NDMA-treated mice 48h post-exposure. The proportion of proliferating cells was determined by flow cytometry of Ki67-positive cells. Data represented as mean ± SD. For all graphs, each data point represents one mouse. Statistical comparisons calculated by unpaired Student’s t-test, *p < 0.05, **p < 0.01, ^#^p < 0.0001.

### Slide 6
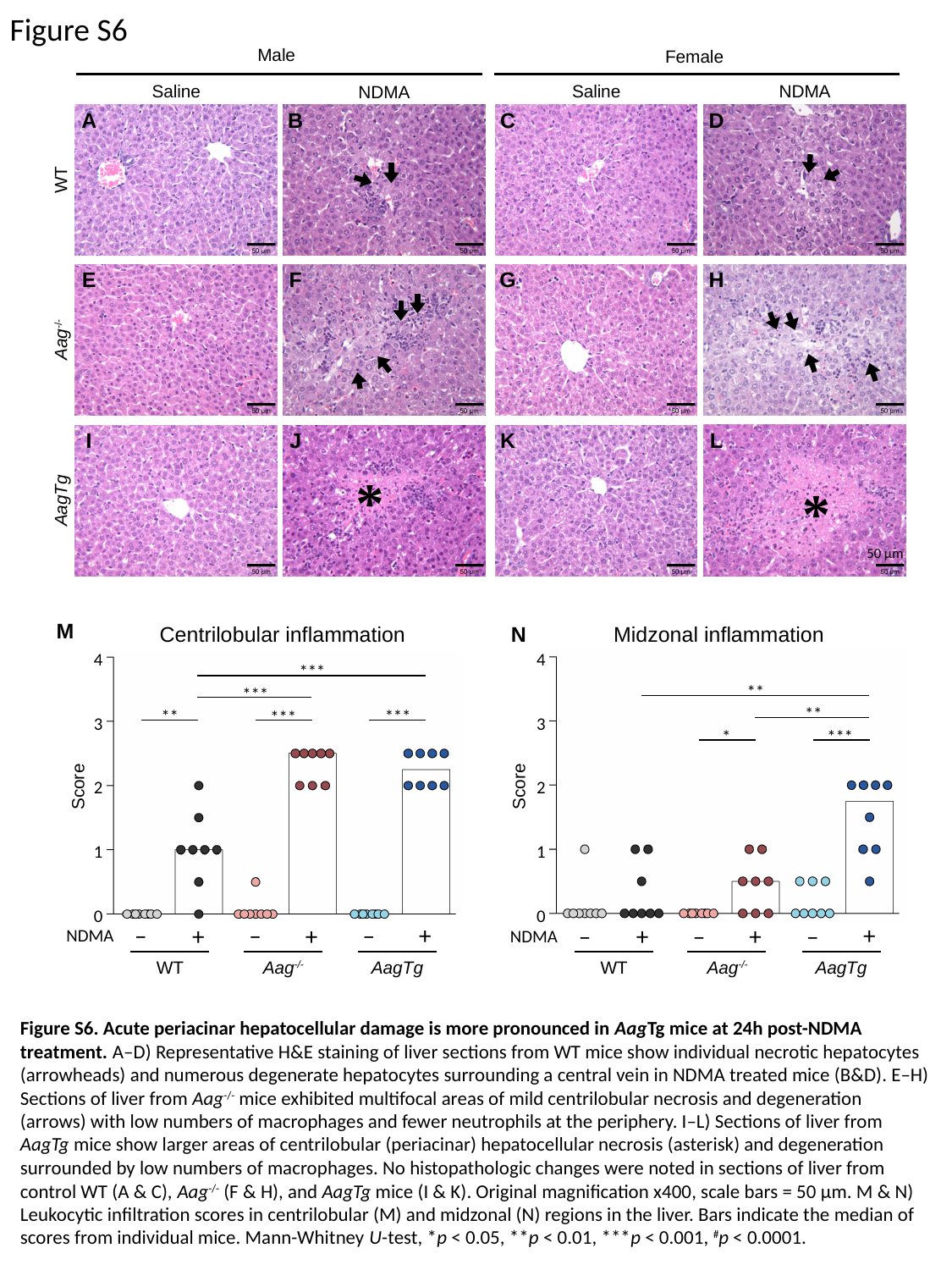

Figure S6
Male
Female
Saline
Saline
NDMA
NDMA
A
B
C
D
WT
E
F
G
H
Aag^-/-^
I
J
K
L
*
*
AagTg
50 μm
M
N
Centrilobular inflammation
4
***
***
**
***
***
3
2
Score
1
0
+
–
–
–
+
+
NDMA
AagTg
Aag^-/-^
WT
Midzonal inflammation
4
**
**
3
***
*
2
Score
1
0
+
–
–
–
+
+
NDMA
AagTg
Aag^-/-^
WT
Figure S6. Acute periacinar hepatocellular damage is more pronounced in AagTg mice at 24h post-NDMA treatment. A–D) Representative H&E staining of liver sections from WT mice show individual necrotic hepatocytes (arrowheads) and numerous degenerate hepatocytes surrounding a central vein in NDMA treated mice (B&D). E–H) Sections of liver from Aag^-/-^ mice exhibited multifocal areas of mild centrilobular necrosis and degeneration (arrows) with low numbers of macrophages and fewer neutrophils at the periphery. I–L) Sections of liver from AagTg mice show larger areas of centrilobular (periacinar) hepatocellular necrosis (asterisk) and degeneration surrounded by low numbers of macrophages. No histopathologic changes were noted in sections of liver from control WT (A & C), Aag^-/-^ (F & H), and AagTg mice (I & K). Original magnification x400, scale bars = 50 μm. M & N) Leukocytic infiltration scores in centrilobular (M) and midzonal (N) regions in the liver. Bars indicate the median of scores from individual mice. Mann-Whitney U-test, *p < 0.05, **p < 0.01, ***p < 0.001, ^#^p < 0.0001.

### Slide 7
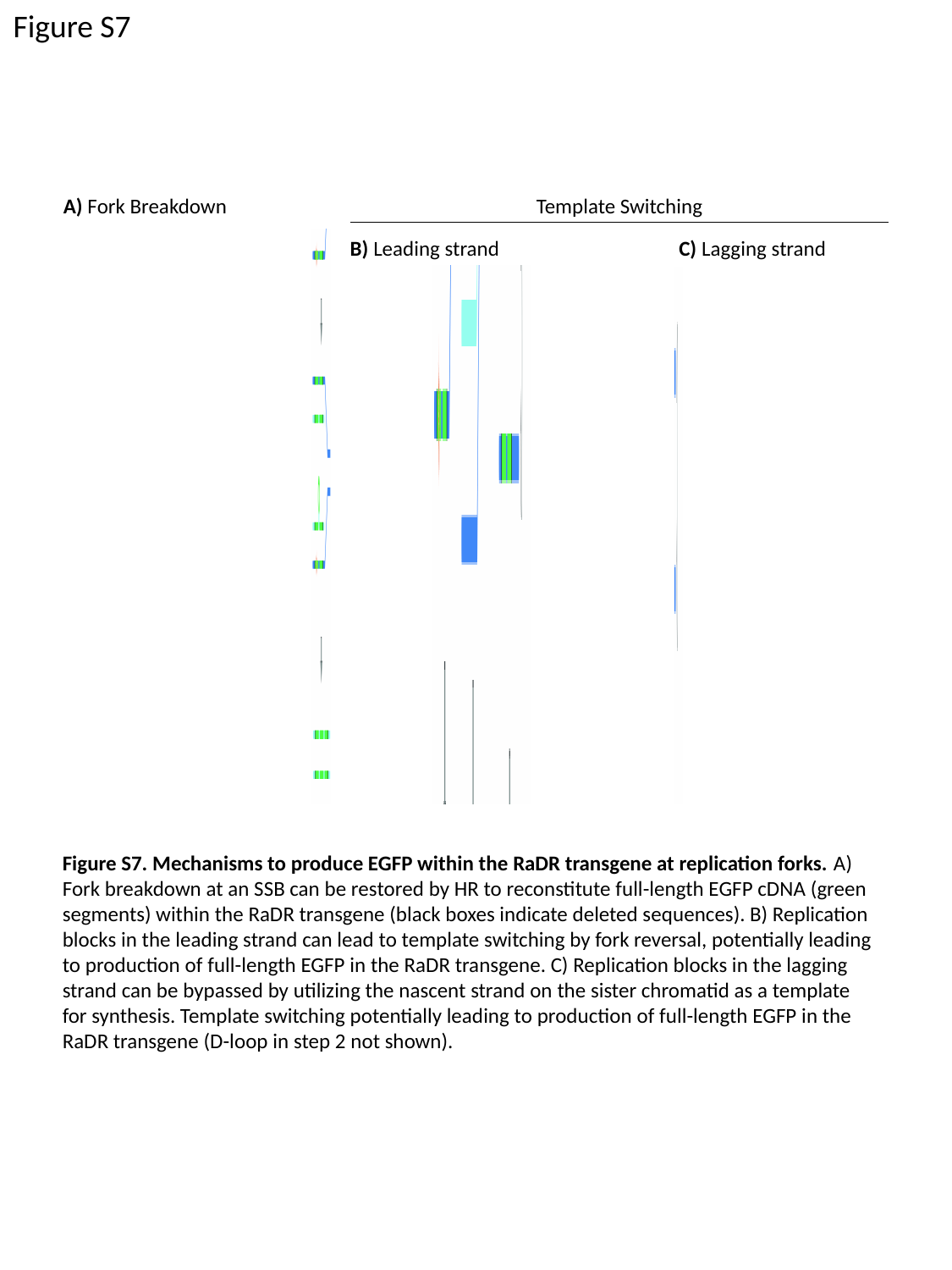

Figure S7
A) Fork Breakdown
Template Switching
B) Leading strand
C) Lagging strand
Figure S7. Mechanisms to produce EGFP within the RaDR transgene at replication forks. A) Fork breakdown at an SSB can be restored by HR to reconstitute full-length EGFP cDNA (green segments) within the RaDR transgene (black boxes indicate deleted sequences). B) Replication blocks in the leading strand can lead to template switching by fork reversal, potentially leading to production of full-length EGFP in the RaDR transgene. C) Replication blocks in the lagging strand can be bypassed by utilizing the nascent strand on the sister chromatid as a template for synthesis. Template switching potentially leading to production of full-length EGFP in the RaDR transgene (D-loop in step 2 not shown).
